# Evaluating SARS-CoV-2 Antibody Resilience via Prediction and Design of Escape Viral Variants

**DOI:** 10.1101/2025.05.12.653592

**Authors:** Marian Huot, Pierre Rosenbaum, Cyril Planchais, Hugo Mouquet, Rémi Monasson, Simona Cocco

## Abstract

The evolutionary trajectory of SARS-CoV-2 is shaped by competing pressures for ACE2 binding, viability, and escape from neutralizing antibodies targeting its receptor-binding domain (RBD). Here, we present EscapeMap, a modular framework that enables prediction and design of variants escaping antibodies. EscapeMap integrates deep mutational scanning data for ACE2 and 31 monoclonal antibodies with a generative sequence model trained on pre-pandemic Coronaviridae. To experimentally probe escape potential, we designed RBD variants under pressure from four clinically relevant antibodies (SA55, S2E12, S309, VIR-7229). Among these designs, bearing up to 21 mutations from wildtype, 50% expressed as stable proteins. Binding assays confirm that S309 and VIR-7229 retain recognition across diverse mutation combinations. EscapeMap accurately forecasts which antibodies are vulnerable to escape by our designed sequences. Finally, by identifying correlated escape routes, we predict and experimentally verify, antibody combinations less prone to simultaneous escape, offering a quantitative basis for guiding therapeutic strategies.

## INTRODUCTION

The capacity of SARS-CoV-2 to acquire mutations that enhance transmissibility, evade humoral immune response, or both, poses a challenge to pandemic preparedness^1^. This evolutionary pressure is particularly evident in the spike protein’s receptor-binding domain (RBD), which not only mediates viral entry but also serves as a key target for neutralizing antibodies^2^. Mutations in this region that reduce antibody neutralization allow the virus to evade immune responses, thereby fueling new pandemic waves. Current strategies to mitigate pandemic waves typically assess vaccine and antibody efficacy against past or circulating variants. This retrospective approach is inherently limited, as it fails to measure robustness against future variants that are most likely to emerge and to continue the cycle. A conceptual shift is necessary to develop strategies to proactively evaluate therapeutics against putative new variants rather than existing ones only.

Computational works, especially evolutionary data-driven approaches^3,4^have proven effective in identifying future mutations. Previous work^5^ demonstrates an advancement in this regard, as it leverages structural information to assess the accessibility of epitopes and the likelihood of mutations disrupting antibody binding without compromising protein stability.

On the other hand, experimental techniques such as deep mutational scanning (DMS) and pseudovirus assays provide high-resolution, functional insights into how mutations impact ACE2 binding and antibody neutralization. DMS systematically introduces mutations across a protein and measures their effects on binding, expression, or function using high-throughput selection assays coupled with deep sequencing. This approach has been applied to SARS-CoV-2 to map escape mutations for multiple antibodies^6–14^, paving the way for a deeper understanding of RBD evolution. As a matter of fact, several studies^15–18^ successfully leveraged experimental data to enhance the prediction of high-fitness SARS-CoV-2 variants, by incorporating DMS or combinatorial scanning results into their models.

Despite extensive mapping of antigenic change, our ability to evaluate the likelihood of viral mutations as a function of specific immune pressures remains limited. Existing approaches either estimate static mutational risk from generic constraints^5^, or predict binding without a generative, selection-tunable model^19,20^. Our study addresses these limitations by developing EscapeMap, a probabilistic framework that explicitly parameterizes the selective environment, by integrating a generative sequence model trained on pre-pandemic evolutionary data with a biophysical model trained on DMS data for ACE2 and antibody binding. EscapeMap’s fitness landscape is not static. Antibody selective pressures can vary over time, enabling a quantitative understanding of viral behavior as a function of immunity. The model’s generative capacity allows us to design artificial, highly mutated, yet viable RBD sequences. We leverage this capability to “stress-test” therapeutic antibodies by computationally designing and then experimentally validating, by binding assays, functional variants engineered to escape them, thereby revealing their potential vulnerabilities to viral evolution.

## RESULTS

### RBD mutational landscape from Deep Mutational Scans and Pre-pandemic sequence data

We developed EscapeMap, a computational model to investigate the landscape governing mutations in the receptor-binding domain (RBD) of SARS-CoV-2, balancing protein viability—including expression, stability and overall functionality— with the capacity to bind to the human host cells, and antibody escape, which is a critical factor driving viral adaptation. The model evaluates the likelihood of a RBD sequence by integrating three main components (Figure 1a): ACE2 affinity, antibody escape, and sequence viability.

**Fig. 1.**
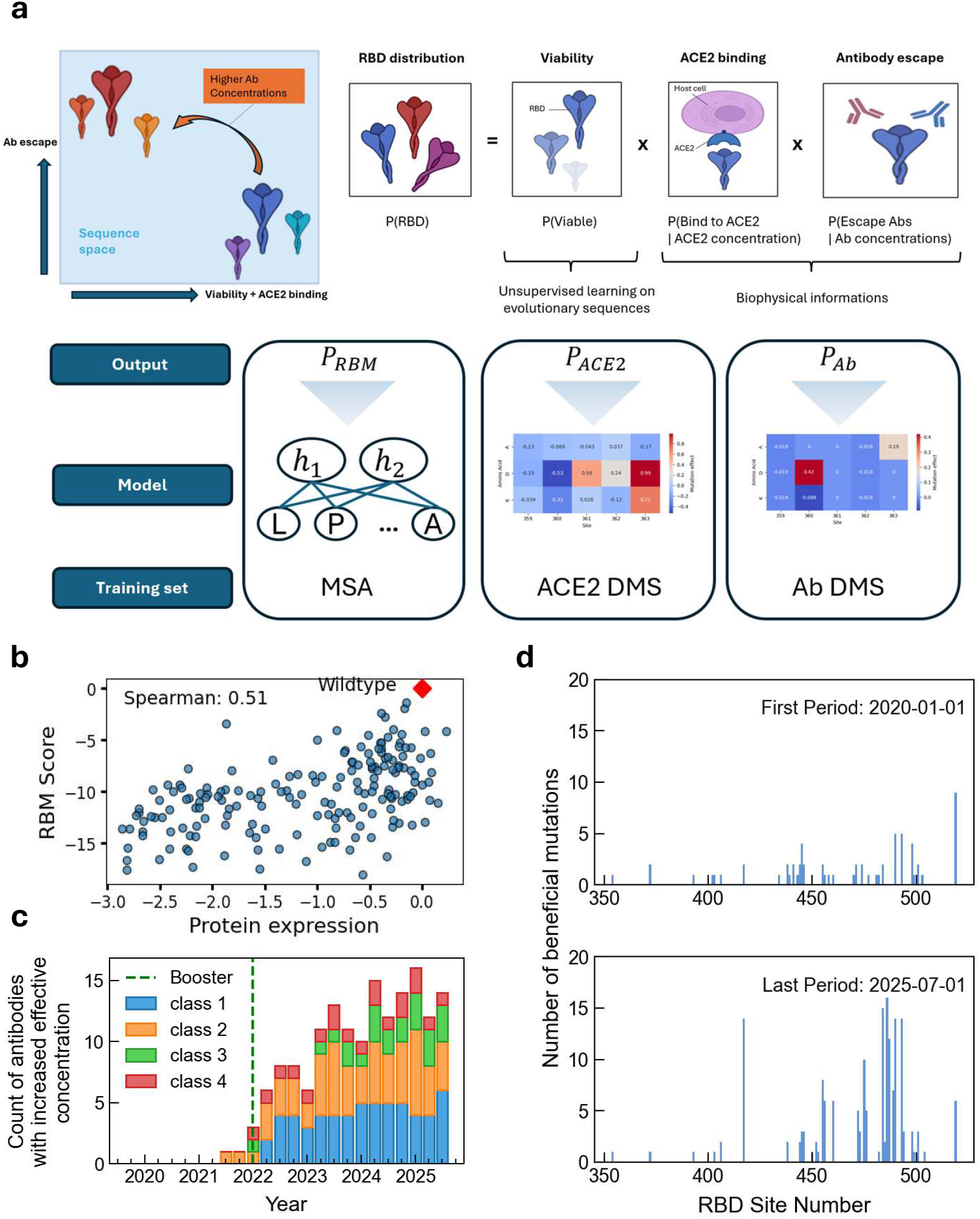
EscapeMap captures shifting mutational landscape and rising immune pressure during the pandemic. (a) Model architecture. Variants with high EscapeMap likelihood must be viable and show good ACE2 binding and antibody escape. Viability is derived from a RBM fitted on a multiplesequence alignment of 2610 prepandemic Coronaviridae sequences. ACE2 binding and antibody escape probabilities are obtained from a biophysical model fitted on deep mutational scans. (b) Scatter plot of RBM mutational scores and experimentally measured changes in expression ^22^ of single-site mutants, averaged per site, in the Wuhan background. (c) Number of antibodies with fitted concentration that increased by a factor of 2 compared to the initial time period, per class and across times. Effective antibody concentrations were fitted on Nextstrain data every 3 months. (d) Number of beneficial mutations per site on the WT background, according to models fitted with parameters from different time periods. Beneficial mutations are defined as single-site variants having a model score greater than the wildtype (WT).

To account for the overall viability constraints for RBD sequences, we learn an unsupervised model from naturally occurring protein sequences which have been selected through evolution. We model the likelihood that a sequence *v* belongs to Coronaviridae RBD protein family, *P*_*RBM*_ (*v*), using a Restricted Boltzmann Machine (RBM), trained on multiple sequence alignments (MSA) of natural prepandemic RBD sequences in the Coronaviridae family, which are not restricted to human Coronavirus only. The RBM is a generative neural network with two layers, capturing latent factors of co-evolution by forming a joint probability landscape over observable sequence states and their hidden representations^21^.

To model ACE2 binding, we introduce the probability *P*_*ACE*2_(*v*) of the free RBD protein with sequence *v* to bind ACE2. The ACE2 binding probability is parametrized by the effective ACE2 concentration, which is treated as a parameter to be fitted from pandemic sequence data, and by the RBD-ACE2 binding free energy. The RBD-ACE2 binding free energy is estimated for each RBD sequence *v* with an additive model (Eq 8) inferred from deep-mutational scans (DMS) data providing mutational effects of single-site mutants of the Wuhan wild-type RBD^22^ (see Methods).

Lastly, the probability of antibody escape captures the selective pressure imposed by the immune system. For each antibody *Ab*_*m*_ considered in the model, the escape probability 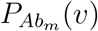 of a RBD variant *v* is parametrized by the binding free energy between RBD and *Ab*_*m*_ and the effective antibody concentration. Similarly to ACE2 binding affinity, the binding free energy between RBD and antibody depends on the underlying RBD sequence and is estimated using DMS data^6–12^. The effective concentration can be tuned to control the probability of escape (see Methods). We first focus our model on a panel of SARS-CoV-2 neutralizing antibodies targeting the RBD and belonging to classes 1 to 4^12^. Specifically, our initial dataset includes 7 class 1, 9 class 2, 5 class 3, and 8 class 4 antibodies. Because these antibodies were discovered at the beginning of the pandemic, they most likely have influenced the evolutionary trajectory of SARS-CoV-2 during its initial spread. In addition, they were extensively characterized in DMS experiments, providing rich quantitative data necessary to estimate binding free energies for every variant (Eq. 8).

Crucially, EscapeMap is modular: while the total likelihood is defined by the product 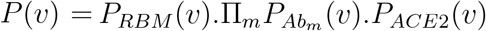, antibody pressure can be excluded, yielding a generative landscape defined by ACE2 binding and RBM-based viability only (ACE2–RBM), or included to account for immune selection via antibody binding probabilities.

### Validation of EscapeMap components

We first extensively benchmark the predictive power of our models for ACE2 and antibody binding (Figure S1). For antibody binding, additive models for the binding free energy fitted on DMS data have previously been shown to perform well for predicting fold changes in neutralization^23^. We further benchmark this component on combinatorial antibody datasets for 3 antibodies that lost binding to BA.1^24^. We obtain very high AUC values (*>* 0.8, Figure S1a), indicating that predictive accuracy remains strong even for multi-mutant variants. We then evaluate how the antibody escape term predicts BA.1 escape on a panel of 438 antibodies identified in WT convalescents^13,14^ and find strong agreement between prediction and experimental data (AUC = 0.73), above the 0.5 baseline (Figure S1b). For ACE2 binding, we use a combinatorial dataset containing all combinations of BA.1 mutations^25^. We observe only a minor loss in predictive power when moving from single to multi-mutant variants. Spearman correlation decreases from *ρ* = 0.72 for single variants to *ρ* = 0.59 for mutants with at least 7 mutations (Figure S1c). We compare our approach to an ACE2-binding predictor from the literature (MLAEP)^20^ and to a zero-shot ESMIF binding predictor^26^; both fail to show predictive power on this dataset, likely because all test sequences are viable and these models cannot capture fine-grained differences in affinity. These results support the use of our DMS-informed additive model of mutational effects for the binding free energy as a robust and interpretable tool for modeling ACE2 binding.

To validate the RBM component trained on an MSA containing data from pre-pandemic coronaviruses sequences only, we first show that the RBM-generated sequences successfully capture the evolutionary pairwise epistatic patterns of the natural data (Spearman *ρ* = 0.57, Figure S2d). Moreover we assess whether the RBM log-likelihood score, log *P*_*RBM*_, serves as a reliable proxy for protein viability, in particular for expression. To do so, we compute a site-dependent RBM mutational score (see Methods), which reflects the average impact of mutations at a site on protein viability. Consistent with experimental findings^3^, we observe a strong correlation between protein expression (measured on single mutants^11^) and the RBM score obtained with our epistatic model (Figure 1b, Spearman’s *ρ* = 0.51), which outperforms a non-epistatic (independent-site) model (*ρ* = 0.24, Figure S2c). Lastly, EscapeMap reproduces the mutational entrenchment observed in ACE2 binding DMS data^27^, showing that BA.1 mutations are overall more favorable in the BA.1 genetic background than in the wild-type background (Figure S3a). The same pattern is observed for the BA.2 background (Figure S3b). Interestingly over the 14 substitutions in BA.1 with respect to the WT, the RBM component alone reproduces the positive background effects for 9 mutations and the negative one for 1 mutation (Q493R, which was reversed in later variants). Similarly on the BA.2 background the sign of the entrenchment is correctly predicted for 73% of the 15 substitutions. The full ACE2-RBM model improves such predictions (Figures S3c-d). Specifically, the N501Y mutation increases the EscapeMap (RBM+ACE2) score on the BA.1 and BA.2 background, thanks to the non-linear transformation used to model ACE2 binding probability. Overall EscapeMap (RBM+ACE2) model correctly predicts the sign of the entrenchment in ACE2 binding DMS for respectively the 92% (BA.1) and 80% (BA.2) of the substitutions. The above findings confirm the importance, for mutations improving the ACE2 bindings, of background stabilizing effects (modeled by the RBM component) combined with additional epistatic effects in key binding sites (modeled by the ACE2 binding probability), as previously observed^22^.

### Increasing immune breadth drives viral mutational exploration

We next investigate how antibody escape contributes to viral evolution during the pandemic, in conjunction with ACE2 binding affinity and viability constraints captured by the RBM. To assess temporal adaptation, we divide the pandemic into three-month non-overlapping intervals spanning the 2020 to 2025 period. For each period, we fit the model parameters to maximize the likelihood of observed sequences (see Methods), yielding effective concentrations for the 29 antibodies considered and for ACE2, and RBM inverse temperature (tuning the selective pressure on viability with respect to those on ACE2 binding and antibody escape). Interestingly, we find that while the ACE2 effective concentration remains stable, the RBM inverse temperature decreases (Figure S4), indicating viability constraints become a smaller driver of RBD evolution. On the other hand, after the widespread administration of the booster dose (third vaccination dose), we observe greater antibody diversity, with a high number of different antibodies —primarily from classes 1 and 2— associated with higher effective concentrations compared to early pandemic (Figure 1c). This is consistent with experimental findings^28^, which documented a broader range of antibodies in donors after the third (SN3) vaccination dose (Figure S5b). The observed increase in fitted antibody effective concentrations suggests a progressive shift in the accessible variant space (Figure S6). Early in the pandemic, most residues showed no selective advantage for mutation, consistent with a small number of mutations in observed variants Alpha and Delta. However, under late-pandemic antibody effective concentration parameters, more mutations become beneficial (Figure 1d).

### Assessing EscapeMap predictive power for viral evolution

We evaluate EscapeMap’s predictive power by fitting it on early pandemic data (second quarter of 2020) and comparing its predictions against mutations observed until 2025. When benchmarked against EVEscape; EscapeMap’s top-decile predictions show a slightly higher success rate in capturing mutations that later became common (observed ≥ 100 times) (Figure 2a). While both models capture more than 40% of all existing mutations within their top decile of predictions, they differ in the types of mutations to which they assign the highest scores. Specifically, EscapeMap prioritizes mutations that appeared with high frequency during the pandemic, compared to EVEscape (Figure 2b), suggesting that EscapeMap is more sensitive to the features that drive widespread viral fitness.

**Fig. 2.**
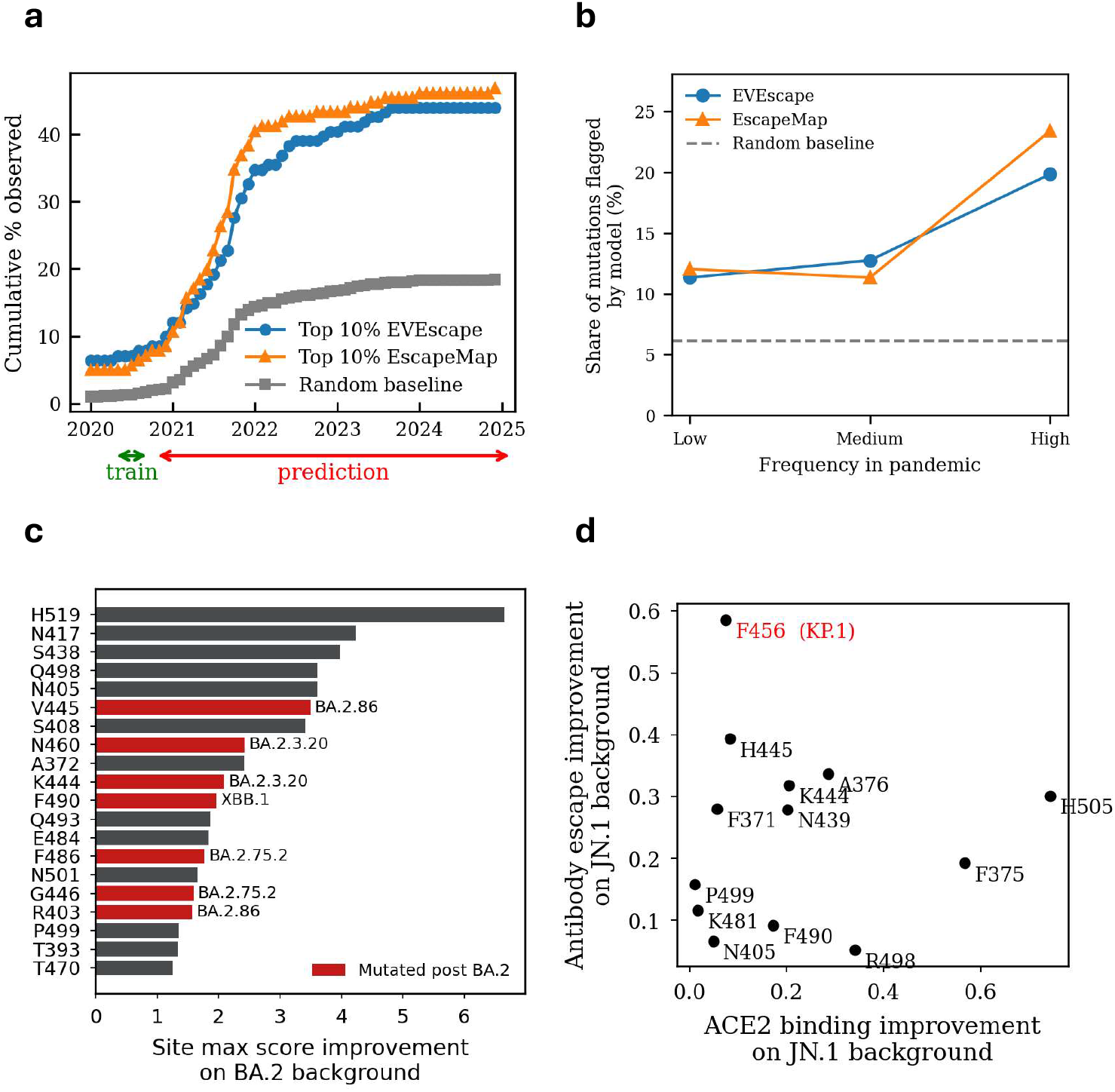
EscapeMap predicts future mutations on different backgrounds. (a) Percentages of top decile predicted escape mutations by EscapeMap (fitted on sequences of second quarter of 2020) and EVEscape, seen more than 100 times by each date since the start of the pandemic. Analysis focuses only on non-synonymous point mutations that are a single nucleotide distance away from the Wuhan wildtype. (b) Percentages of observed pandemic mutations, categorized by observed frequency during the pandemic, within model top score decile. Mutations are stratified into three equal-size bins based on their occurrences: Low (occurrence ≤ 284), Medium (284 *<* occurrence *<* 1443), and High (occurrence ≥ 1443). High-frequency mutations, in particular, are well captured by EscapeMap. (c) Top 20 BA.2–background sites by EscapeMap max–score improvement, using the model fitted during the early BA.2 period (first quarter of 2022) and expanded with antibodies from BA.1-convalescent individuals ^13,14^. Red bars indicate sites that later mutated in BA.2 sublineages with annotations marking the first sublineage where each mutation appeared. (d) Sites that improve antibody escape while maintaining ACE2 binding and viability. Shown are top-ranked mutations on the JN.1 background from the model fitted on third quarter of 2023. Red labels correspond to post-JN.1 VOC mutations.

Beyond this retrospective validation, we test the model’s ability to forecast the evolution of specific variants. To predict site mutability on BA.2 background, we fit an expanded model to the BA.2 period (first quarter of 2022), with 671 additional antibodies isolated from BA.1-convalescent individuals^13,14^. This expanded model, which includes 700 antibodies compared to the 29 in the original model, better predicts the most widespread, high-frequency mutations that emerged post-BA.2, even if the original Ab panel already captured the main escape pathways (Figure S7). In particular, EscapeMap’s top 20 predicted escape sites (Figure 2c) accurately highlight sites that subsequently mutated in emerging BA.2 sublineages. We finally use the expanded model, fitted in the fourth quarter of 2023 to predict the mutations on the JN.1 background (Figure 2d). Within model–identified sites of interest, we find site 456 as a key candidate for synergistic improvement, coherent with previous experimental studies^29,30^, which also highlighted this region’s importance for enhancing both antibody evasion and ACE2 binding. This site was later mutated in KP.1 and its descendants. Notably, the model consistently captures 50% of the emerging mutational sites across both BA.2 and JN.1 lineages (7*/*14 and 1*/*2 respectively). The presence of only one observed mutation in JN.1 background can be due to a narrower evolutionary window rather than to a reduction in predictive performance.

Lastly, to evaluate the added value of the RBM at different time points, we fit the model on a given three-month window (training set) and evaluate whether the model score is predictive of what is observed in the next three-month period (test set). We consider two tasks: the appearance of a specific mutation at least 100 times, and the identification of sites that developed at least one mutant with at least 100 appearances during the period. Performance is measured using the Area Under the Curve (AUC), which quantifies the model’s ability to correctly identify these true positive mutations and sites against the rate of incorrectly identified false positives across all classification thresholds. Incorporating the RBM term in the EscapeMap model improves predictive accuracy for both the prediction of single-amino-acid variant effects (AUC of 0.81 with RBM vs. 0.77 without RBM) and site-level mutational tolerance (AUC of 0.77 with RBM vs. 0.74 without RBM) compared to simpler models that consider only ACE2 and antibody binding components^16,18^, as shown in Figure S8.

### Generation of artificial variants

The ability to predict likely mutations should translate into the capacity to generate viable viral sequences, enabling the evaluation of treatment efficacy and vaccine resilience across diverse viral evolutionary scenarios.

To do so, we evaluate the structural consistency of the mutants generated by Monte-Carlo sampling of the model at varying the overall inverse temperature parameter *β* (see Methods)^31^, using ESM Inverse Folding (ESM-IF)^26^ scores. ESM-IF is a deep learning model trained on millions of protein structures that estimates the likelihood of a given sequence folding into a specific structural template. Raising the overall inverse temperature parameter *β* (see Methods) strengthens the selective pressures across all components of the model, biasing the sampling towards sequences that better satisfy structural constraints and maintain the binding of ACE2, thus favoring functional and evolutionarily plausible variants (Figure 3a). We then explore the influence of antibody selection pressures on the generated sequences (see SI Figure S9b). Mutants generated using EscapeMap with Ab selection pressure of one antibody exhibit ESMIF scores comparable to those generated without antibody pressure, indicating that the overall structural and functional integrity of sequences is preserved across different selection regimes. Furthermore, we compare the site-wise entropy (see Methods) of our generated sequences with data from the pandemic, finding a positive correlation (Spearman *ρ*=0.62, Figure S9a), although the generated sequences generally exhibited higher variability. This diversity stems from the RBM component being trained on homologous RBD sequences, including those evolutionarily distant from the wild type, while pandemic sequences show higher conservation due to shorter evolutionary timescales. Notably, several sites associated with escape from multiple antibodies—among the 29 antibodies studied—show substantial mutational tolerance, with some sites accommodating over four viable amino acids (Figure 3b).

**Fig. 3.**
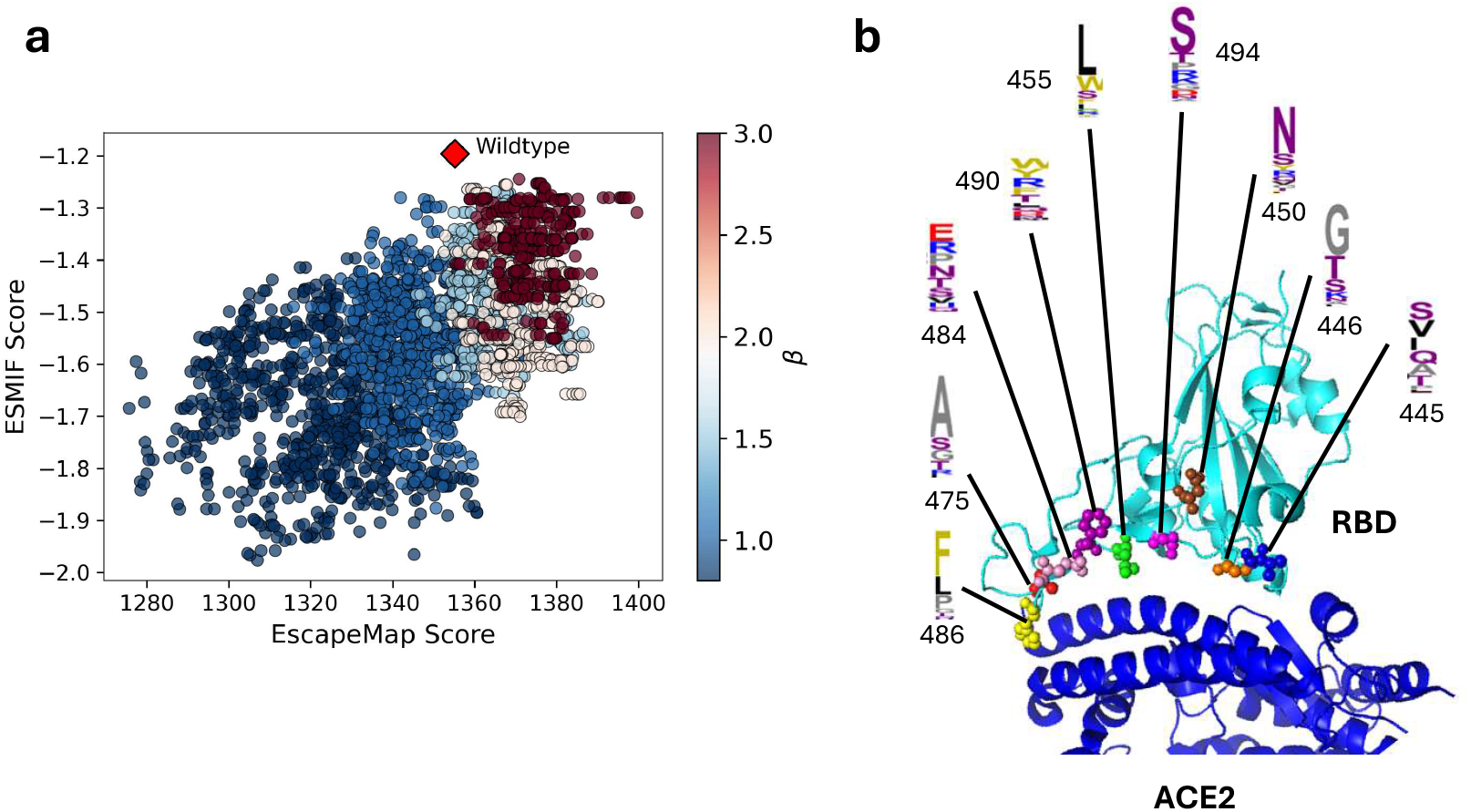
Validation of generative model and structural mapping of high-entropy escape hotspots. (a) Scatter plot of ESM Inverse Folding (ESMIF) score versus EscapeMap score (ACE2–RBM) for generated variants at inverse temperatures *β* ∈ {1, 1.5, 2, 3} without antibody pressure. (b) Structural representation of the RBD (light blue) in complex with ACE2 (dark blue) (PDB: 6M0J). Colored spheres indicate sites involved in escape to at least 3 antibodies, and with entropy above 1. Amino acids show representative mutations at each site.

### Evaluation of antibody cocktails

To quantify the potential for antibody escape, we first use our generative model to sample a large ensemble of functional RBD sequences, defined as those with high RBM and ACE2-binding likelihoods. We then evaluate this *in silico* library against the escape profile of each antibody. This process allows us to compute a key metric: the model escape fraction (*F*_*m*_). This score represents the proportion of all viable, generated sequences predicted to escape a given antibody. A higher *F*_*m*_ score indicates that a large number of functional escape pathways exist, signifying a greater probability that the antibody will rapidly lose efficacy. Applying this framework, we find clear differences in antibody vulnerability. As illustrated in Figure 4a, LY-CoV016 exhibits a high *F*_*m*_ compared to S309^32^, indicating that LY-CoV016 is far more susceptible to immune escape.

**Fig. 4.**
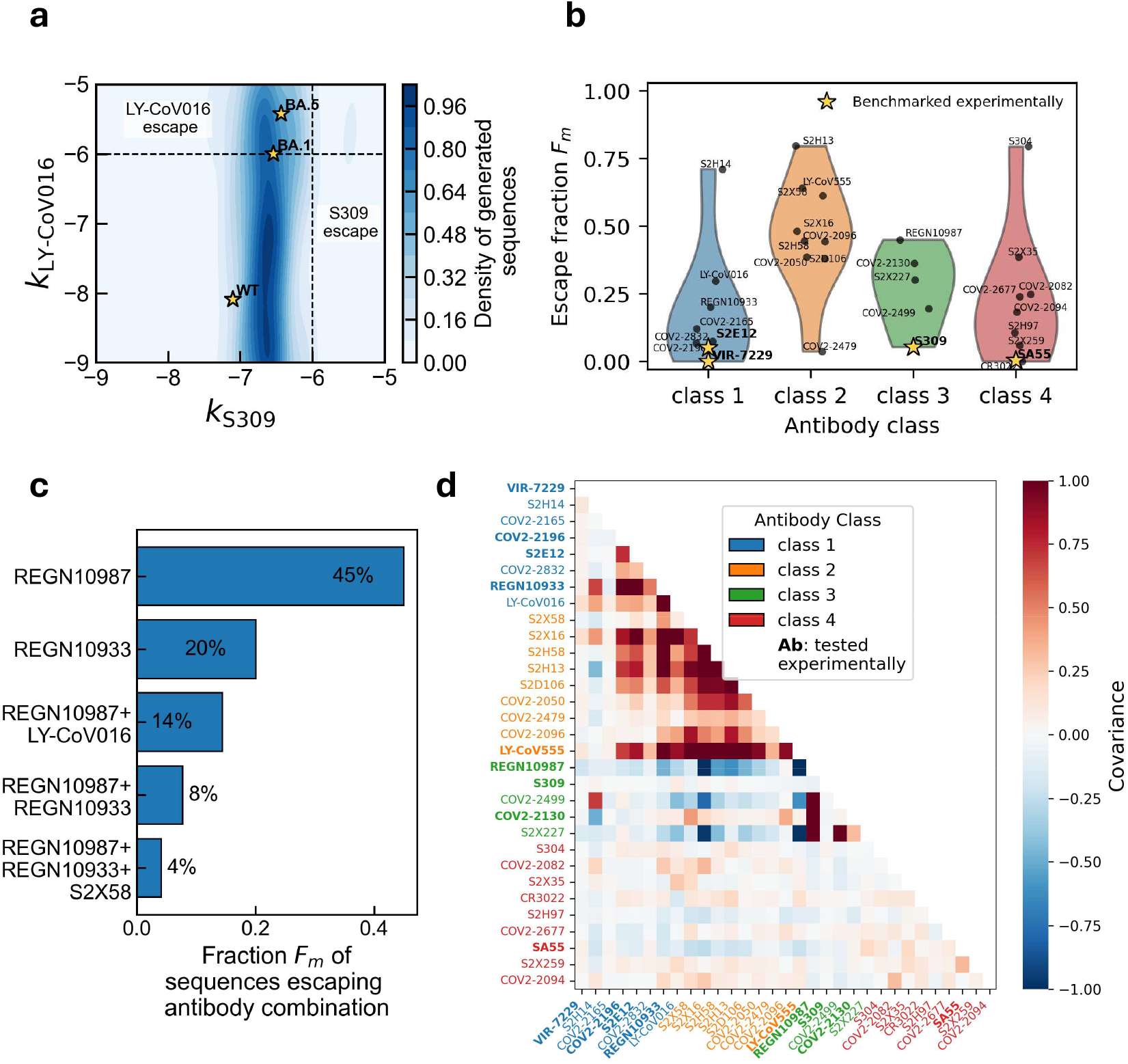
Fraction of escaping sequences informs antibody cocktail design. (a) Predicted binding landscape of generated RBD variants. Dashed lines mark escape thresholds; stars denote VOCs. Variants in the upper-left region are predicted to escape LY-CoV016 while retaining S309 binding. (b) Computed model escape fraction *F*_*m*_ values for each antibody, grouped by antibody class. Antibodies with low escape fractions—SA55, VIR-7229, S2E12, and S309—are highlighted and selected for binding assay. (c) Escape fraction by antibody combination. Bars show the fractions of generated sequences predicted to escape all antibodies in each combination. (d) Model covariances between antibody binding freeenergies. Red indicates positive covariance; blue indicates negative covariance. Experimentally validated covariations are highlighted in bold.

We compute the escaping fraction *F*_*m*_ for the 29 antibodies considered up to now, and for two recently characterized, broad, and potent neutralizers, SA55 and VIR-7229^33,34^. Among the antibodies with the highest resilience (i.e., lowest *F*_*m*_), we identify VIR-7229, S2E12, S309 and SA55 (Figure 4b). These antibodies are selected for experimental benchmarking in the later stages of this study. Interestingly, evolutionary constraints imposed by RBD viability and ACE2 binding substantially limit the overall potential for antibody escape. The proportion of functional escape variants (sampled under model distribution) is lower than that predicted under a uniform sequence distribution (Figure S10a).

Next, we demonstrate how EscapeMap can be used to evaluate the design of effective antibody cocktails. A crucial factor in cocktail efficacy is whether the antibodies have overlapping escape profiles; if they target similar epitopes, a single viral mutation might confer broad resistance across the entire cocktail^35–37^. Our framework allows us to directly quantify such effect. We extend our escape fraction metric to combinations, calculating the proportion of viable, generated sequences predicted to escape all antibodies in a given cocktail (Figure 4c). As expected, adding antibodies to a cocktail tends to decrease the *F*_*m*_ score, as fewer viable sequences possess the mutations required to evade all antibodies simultaneously. However, this decrease is most important when the antibodies are de-correlated, like REGN1087 and REGN10933. We quantify antibody synergy for each pair of antibodies in the panel by computing the covariance between the binding free-energies to the two antibodies of EscapeMap-generated RBD sequences (Methods). As shown in Figure 4d, Antibody cocktails combining class 1 and class 2 antibodies are suboptimal due to their positive covariances, which make them susceptible to shared escape mutations. In contrast, class 3 antibodies display decorrelated or anticorrelated escape profiles when paired with class 1 or class 2 antibodies, reducing the likelihood of shared escape mutations and thereby improving cocktail synergy. This result aligns with experimental findings ^10^, which demonstrated the lack of resilience of the LY-CoV555+LY-CoV016 antibody cocktail—consistent with our EscapeMap’s prediction of a positive covariance. In contrast, the REGN10933+REGN10987 combination, predicted by EscapeMap to have a negative covariance, appears more promising as was experimentally validated ^35^. The concept of escape profile covariance extends beyond pairs to cocktails with three or more Abs (Methods), and can be used to find adequate antibody to add to a pair cocktail such as LY-CoV555+LY-CoV016 (Figure S10b).

### Viral sequence design and in vitro test

To experimentally validate EscapeMap’s generative power, we generate 6 sequences from the distribution without antibody pressure (ACE2-RBM model) at inverse temperature *β* = 3, as well as 16 sequences under selective pressure from VIR-7229, SA55, S2E12 and S309 (ACE2-Ab-RBM model), which demonstrated strong resilience in EscapeMap’s predictions. To generate sequences under high selective pressure, we fix the antibody effective concentrations to a high value *C* = 10^−6^ g/mL in the model probability *P*_*Ab*_ (see Methods).

We achieve a 50% success rate among the tested sequences when evaluating in vitro production of the designed variants (Table 1), a remarkable hit rate given the mutation burden. We attribute this high success rate to the fact that over one-third of the mutations in our designed sequences had already appeared more than 100 times in circulating pandemic variants (Figure S11). In other words, the model successfully captured viable mutations that emerged naturally, which explains the high experimental hit rate. As a control, we also generate 4 sequences with either reduced importance of RBM log-likelihood or ACE2 affinity (see Methods). None of these control sequences could be expressed as recombinant proteins, which emphasizes the importance of maintaining viability and binding potential.

**Table 1:** Number of sequences tested and expressed under various experimental conditions.

| Condition | ACE2-RBM | S2E12 | S309 | SA55 | VIR-7229 | Neg. Ctr. ACE2 | Neg. Ctr. RBM | WT |
| --- | --- | --- | --- | --- | --- | --- | --- | --- |
| Number of sequences tested | 6 | 4 | 4 | 4 | 4 | 2 | 2 | 1 |
| Number of sequences expressed | 3 | 3 | 3 | 1 | 1 | 0 | 0 | 1 |

Interestingly, most functional sequences show good ACE2 binding (Figures 5a-b), although they span a broad range of mutations across the RBD, particularly at the ACE2 binding interface (Figure 5c for structural representation of RBD S2E12c, see Figure S12 for representation of the other sequences that could be expressed). We observe nearly no correlation between ACE2 binding affinity and distance from the wild-type sequence, indicating that our model is capable of generating sequences that are both functionally viable and evolutionarily distant from the wildtype (see Figure S13).

**Fig. 5.**
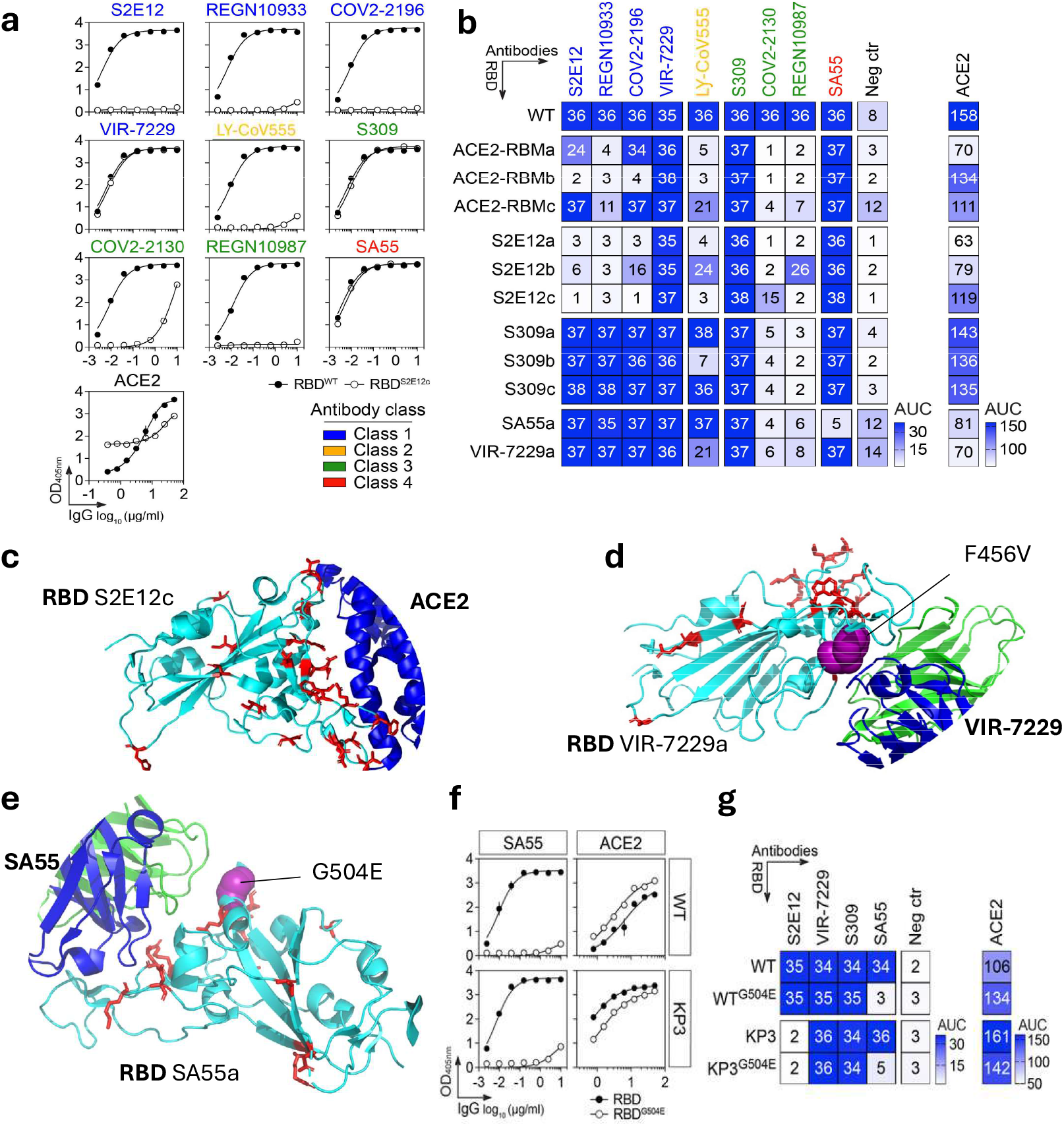
In vitro test of antibody binding on well-expressed designed variants. (a) Representative ELISA graphs showing the binding of anti-RBD antibodies and ACE2 to purified wildtype RBD and S2E12c RBD generated with S2E12 immune pressure. (b) Heatmap of ELISA binding of anti-RBD antibodies and ACE2 to purified designed RBDs (higher scores = higher binding). Tested RBDs: wildtype, 3 designed with ACE2–RBM (no antibody),8 designed with ACE2–Ab-RBM (*c* = − 6 for the Ab indicated in the name of the variants). (c) Structural representation of the RBD (light blue) in complex with ACE2 (dark blue) (PDB: 6M0J). Red highlights indicate sites with a mutation in the tested and expressed variant S2E12c. (d) Structural representation of RBD in complex with ACE2 and the VIR-7229 antibody (PDB: 9ASD). Red = mutations in functional sequences; purple spheres = residue 456 (critical for binding). (e) As in (d), for the SA55 antibody (PDB: 7Y0W); purple spheres = residue 504 (critical for binding). (f) Representative ELISA graphs showing the binding of antibody SA55 and ACE2 to purified wildtype, new variant KP.3, and their G504E mutants. (g) Heatmap of ELISA binding of anti-RBD antibodies and ACE2 to purified wildtype, new variant KP.3, and their G504E mutants.

We experimentally test the Abs binding for the 12 expressed RBD proteins for a panel of 9 antibodies encompassing the 4 antibodies (VIR-7229, SA55, S2E12 and S309) used in the design of the artificial variants under antibody selection pressure (Figures 5a-b). VIR-7229, in particular, proves to be highly resilient, as it retains binding even when residue 456—previously identified as important for binding and already mutated in EG.5 and KP variants—is mutated (Figure 5d).

The escape mechanism observed for SA55 involves a mutation at residue G504E (Figure 5e). Notably, this mutation remains viable even in highly mutated RBD backgrounds, indicating substantial tolerance. To assess whether this mutation alone can drive escape, we introduce G504E into both the wild-type and KP.3 backgrounds. In both cases, the resulting variants maintain strong ACE2 binding while consistently escaping SA55 neutralization (Figures 5f-g). This result demonstrates that G504E is a potent single-point escape mutation for SA55, and that its effect is not dependent on a specific genetic context. The fact that this escape occurs across divergent backgrounds, including the clinically relevant KP.3 variant, reveals a critical vulnerability in SA55’s resilience. To further dissect the specificity of this escape route, we evaluate whether G504E confers escape from other antibodies. We observe no loss of binding to S309 or VIR7229—neither in the WT nor KP.3 backgrounds—indicating that G504E selectively compromises SA55 binding without affecting other antibodies. Together, these results suggest that G504E constitutes a viable and highly specific escape route for SA55, underscoring the importance of considering such context-independent mutations when evaluating antibody robustness.

### Experimental Validation of EscapeMap Escape Predictions

Lastly, we compare model predictions against experimental measurements of binding and escape. We find that EscapeMap’s predictions of antibody escape events were highly accurate, achieving an Area Under the Curve (AUC) of 0.91 (Figure 6a; see SI Figure S13c for per-antibody metrics). The escape threshold *k* ≥ −6 used in our model represents a minimum criterion. This interpretation is consistent with the low false-positive rate ( ∼ 1%) observed in our experimental benchmark while still identifying more than half of the true escape cases. Under this threshold, the model correctly predicts that none of the generated variants escape VIR-7229 (Figure 6b), while COV2-2130 is escaped by most of the expressed sequences.

**Fig. 6.**
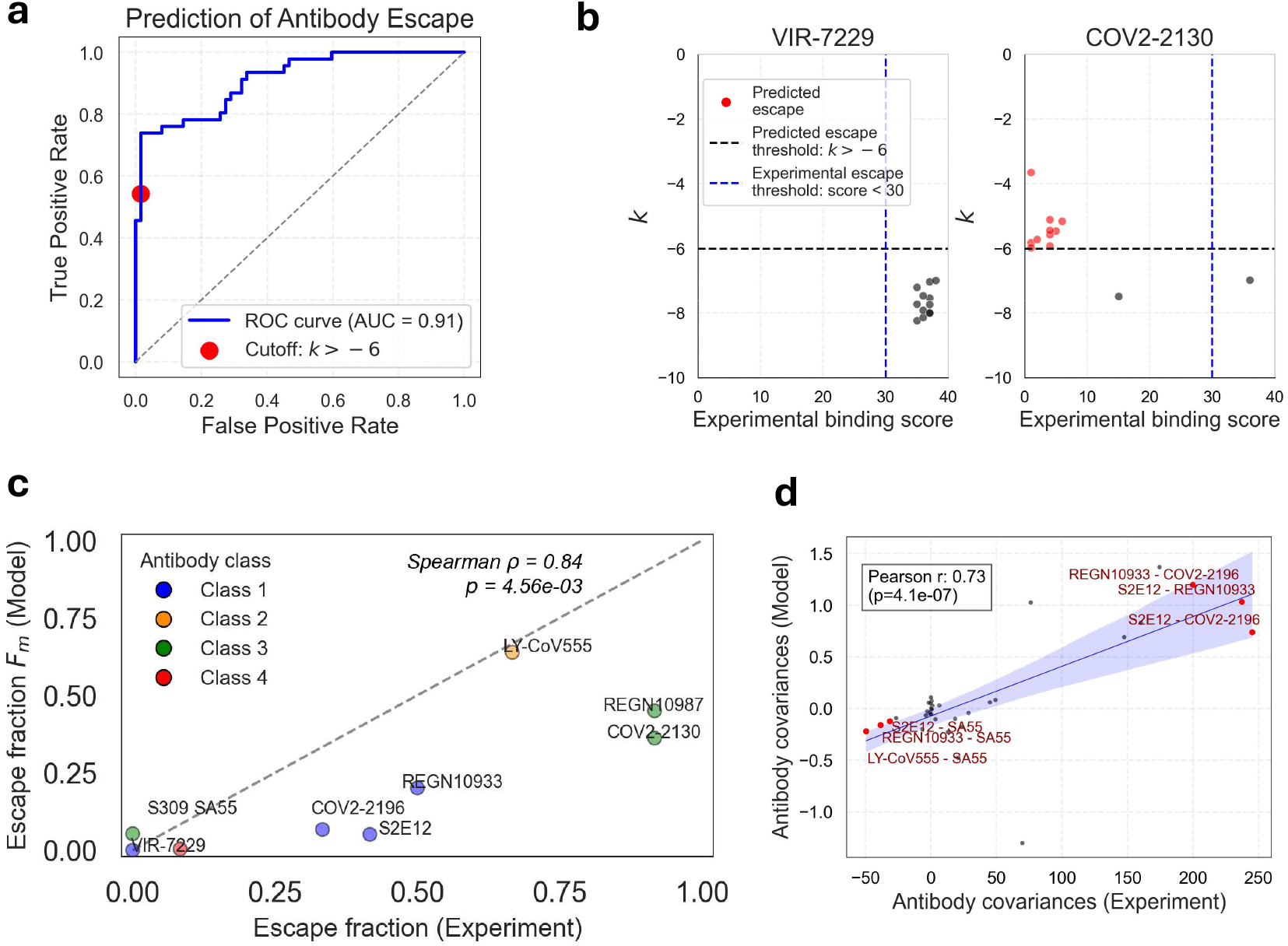
EscapeMap predicts observed antibody escape profiles and antibodies binding covariation. (a) ROC curve for model prediction of antibody escape (experimental binding score *<* 30) on the 11 well-expressed designed variants. (b) Comparison of measured and predicted binding scores for antibodies VIR-7229 and COV2-2130. Each point corresponds to a designed sequence that could be expressed. (c) Percentage of expressed designed sequences escaping each antibody in experiment, compared to model-derived fraction *F*_*m*_ of sequences escaping antibody. (d) Covariance between antibody binding free energies by experiment vs. model. Pairs with highest/lowest experimental covariances are highlighted.

When comparing the escape fraction in experiments and the one computed by the model on a set of 1000 artificial sequences, we find great correlation in the ranking of antibodies (spearman *ρ* = 0.84). Experimental results confirm in particular the existence of multiple sequences capable of escaping class 1 REGN10933^38^ and most of class 2 and 3 antibodies; no RBD was able to escape class 3 S309 and class 1 VIR-7229. The model’s estimated escape fraction tends to be smaller than that observed in experiments, as seen with S2E12 (Figure 6c). This is consistent with our use of a stringent escape threshold as a minimum criterion for high-certainty escape, which prioritizes a low false-positive rate. Additionally, epistatic interactions, which are not included in the additive binding free energy model, likely decrease binding further in these highly mutated sequences. Last of all, we show in Figure 6d that the predicted covariances between antibody binding free-energies match those in experiments on the generated sequences (Pearson correlation: 0.73). Notably, class 1 antibodies systematically show the same binding behavior towards functional generated sequences. We find, both in the model and in the experimental data, that SA55+REGN10933 is a good cocktail as their experimental binding scores are anti-correlated, despite REGN10933 not being a powerful antibody alone. Overall, these results provide a robust experimental validation of our model, demonstrating its potential for predicting viral escape and guiding the design of more resilient antibody therapeutics.

## DISCUSSION

In this study, we develop a generative framework, EscapeMap, to model the evolution of the SARS-CoV-2 receptor-binding domain (RBD) under immune and receptor binding constraints. The model combines three complementary components (sequence viability, ACE2 affinity and antibody escape) and demonstrates several advantages in the context of variant predictions. First, EscapeMap prospectively identifies mutation-prone sites under realistic selective pressures and complements existing approaches. In retrospective evaluations, it anticipates future mutations up to 5 years ahead with strong enrichment over random and a modest gain over EVEscape^5^. Unlike with EVEscape model, EscapeMap’s fitness landscape is not static and explicitly parameterizes the selective environment by fitting antibody-specific and ACE2 effective concentrations, recovering the increasing number of antibodies exerting pressure over time. Post-booster landscape is characterized by a “mutational explosion” driven by intense and diverse antibody pressure. As viability constraints become a secondary driver, the accessible variant space expands, making a wider array of mutations beneficial for viral fitness. This suggests that the virus is not merely drifting but is being actively pushed into new regions of the mutational landscape by the breadth of human immunity, and aligns with research suggesting that mass vaccination may accelerate viral evolution^39^. Crucially, variants are scored according to their loglikelihoods, tailored to identify biologically plausible, high-fitness candidates for surveillance and stress-testing, rather than to predict takeover^17^ and immunity based lineage prevalence^40^. Furthermore, while related in spirit to UniBind^19^ in leveraging ACE2 and antibody binding predictors with a simpler DMS-fitted architecture, EscapeMap differs in how these signals are used: rather than standalone effect predictors, binding contributions enter the generative likelihood, are tunable via antibody effective concentrations, and couple with a viability term to produce thousands of artificial sequences that can be used to stress-test therapeutics.

Second, we introduce a quantitative framework for evaluating antibody resilience against viral evolution. Existing metrics of antibody breadth typically rely on neutralization against circulating variants or structural epitope mapping^9,41^. In contrast, antibody resilience estimates the probability that viable, ACE2-binding sequences can escape a given antibody. This probabilistic approach allows for a systematic ranking of antibodies and for the identification of vulnerable epitopes before escape mutations are experimentally observed. Moreover, by modeling the joint mutational landscape across antibodies, EscapeMap can evaluate antibody cocktails in the presence of epistatic constraints, favoring cocktails for which jointly escaping mutations decrease viral fitness score. This extends beyond traditional cocktail design strategies that focus solely on minimizing epitope overlap^35–37^.

Third, EscapeMap’s generative capacity enables the design of highly mutated yet viable RBD variants^42^. Among 22 designed sequences—each carrying up to 25 substitutions relative to Wuhan-Hu-1—50% expressed successfully, a striking outcome given prior work on the same protein showing expression drops sharply with mutational load, with hit rates falling below 2% beyond eight random mutations^22^. This indicates that EscapeMap does not merely interpolate around known variants but can identify functional configurations in remote regions of sequence space. Mechanistically, the model achieves this by capturing the underlying epistatic landscape. An RBM trained on pre-pandemic sequences learns the higher-order co-evolutionary couplings on the RBD amino-acids essential for structural and functional compatibility, while a two-state sigmoidal model maps ACE2 and antibody contributions to fitness^43,44^. The generative capacity to design a library of viable and highly diverse antigens is directly actionable for vaccine design. For instance, advanced strategies like mosaic nanoparticles^45^ depend on co-displaying a diverse array of RBDs. EscapeMap provides a rational method for generating these components, enabling the design of vaccines that can focus immunity on conserved epitopes and broaden sarbecovirus coverage.

A primary limitation of EscapeMap is its exclusive reliance on sequence data and a prepandemic RBM training set that lacks SARS-CoV-2-specific ACE2 selective pressures; additionally the ACE2 binding affinity is only fitted on DMS experiments on single mutants on the Wuhan background. While the above limitations ensures interpretability in a truly prospective evaluation that does not see SARS-CoV-2 evolution in advance, additional structural and ACE2binding constraints could be integrated thanks to new DMS data^27,46^ and data augmentation thanks to deep learning model such as ESM Inverse folding^26,47^. More generally, as shown by the expanded model fitted on 671 additional antibodies isolated from BA.1 convalescent individuals (Figure 2),^13,14^ the different modules of EscapeMap could be easily updated as new data become available.

## RESOURCE AVAILABILITY

### Lead contact

Further information and requests for resources and reagents should be directed to and will be fulfilled by the Lead Contact, Simona Cocco.

### Materials Availability

This study did not generate new materials.

### Data and code availability

- This paper analyzes existing, publicly available data. These accession numbers for the datasets are listed in the key resources table.
- All original code has been deposited at https://github.com/m-huot/ESCAPE_MAP^48^ and is publicly available as of the date of publication. DOIs are listed in the key resources table.
- Any additional information required to reanalyze the data reported in this paper is available from the lead contact upon request.

## ACKNOWLEDGMENTS

We acknowledge funding from the Agence Nationale de la Recherche (ANR-19 Decrypted CE30-0021-01 to S.C. and R.M.). H.M. received core grant from the Institut Pasteur. P.R. was supported by a two-years postdoctoral fellowship from the Pasteur-Roux-Cantarini program (Institut Pasteur). We thank Dianzhuo Wang for fruitful discussions.

## AUTHOR CONTRIBUTIONS

M.H., H.M., R.M., and S.C. conceived the project, supervised the research, and wrote the manuscript. M.H., R.M., and S.C. developed the computational model and performed data analysis. P.R., C.P. and H.M. designed and carried out the experimental and biological validation.

## Supplementary information

**Fig. S1.**
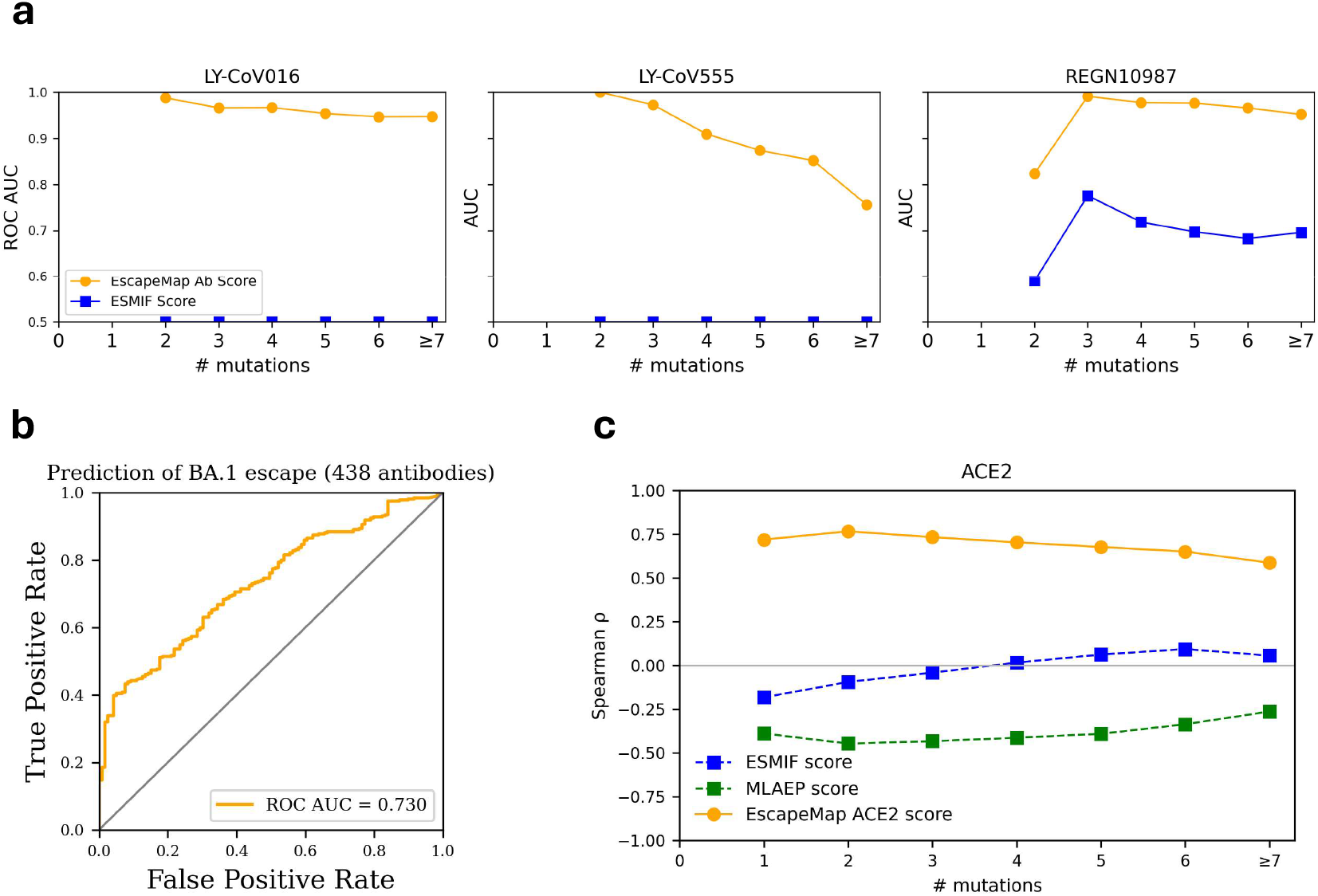
Validation of ACE2 binding and antibody escape modeling. (a) Area under the ROC curve (AUC) for predicting antibody escape in a combinatorial dataset ^24^, shown as a function of mutational load. We compare the EscapeMap antibody score (fitted on DMS data) with the deep learning–based ESMIF ^26^ score. AUC values below 0.5 (random) were put at a 0.5 value. (b) AUC for predicting BA.1 escape ^13,14^ across 438 antibodies isolated from wild-type convalescent samples. (c) Spearman correlation between the EscapeMap ACE2 score and experimentally measured dissociation constants on a combinatorial dataset ^25^. Comparison is made with ESMIF ^26^ and MLAEP ^20^ predictions.

**Fig. S2.**
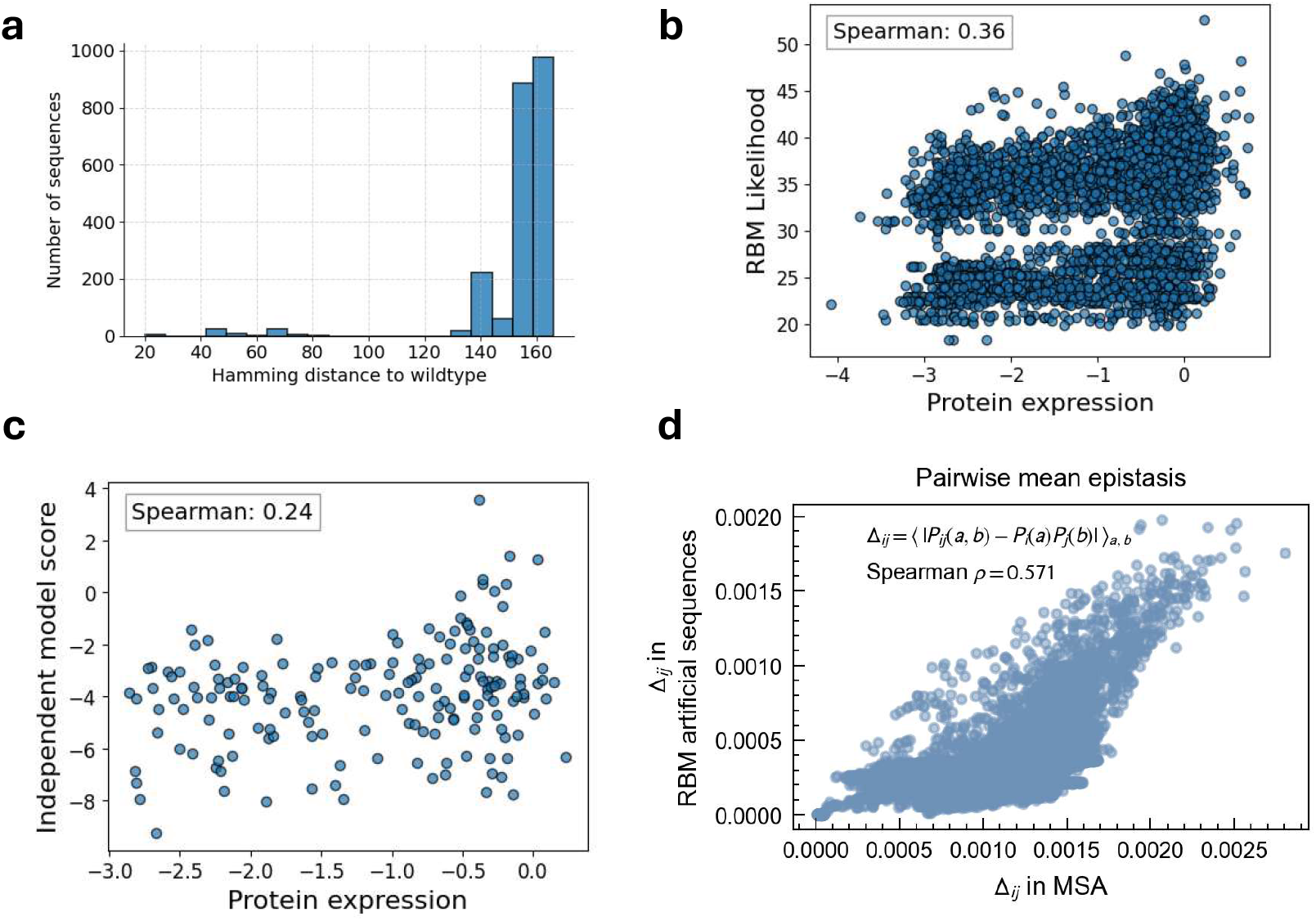
RBM training. (a) Distances between RBD-related sequences in the MSA used to fit the RBM and wildtype RBD (Wuhan). (b) Correlation between RBM mutability scores and experimentally measured expression changes for all single variants on Wuhan background. (c) Correlation between independent-model mutability score and experimentally measured expression changes, averaged per site on Wuhan background. (d) Comparison of pairwise epistasis strength between natural and RBM-generated sequences. Each point represents one residue pair. The *y*-axis shows the mean absolute epistasis Δ_*ij*_ computed from artificial sequences generated by the RBM model, and the *x*-axis shows the corresponding value from the natural multiple sequence alignment (MSA). The RBM reproduces the co-evolutionary coupling structure observed in natural data (Spearman *ρ* = 0.57).

**Fig. S3.**
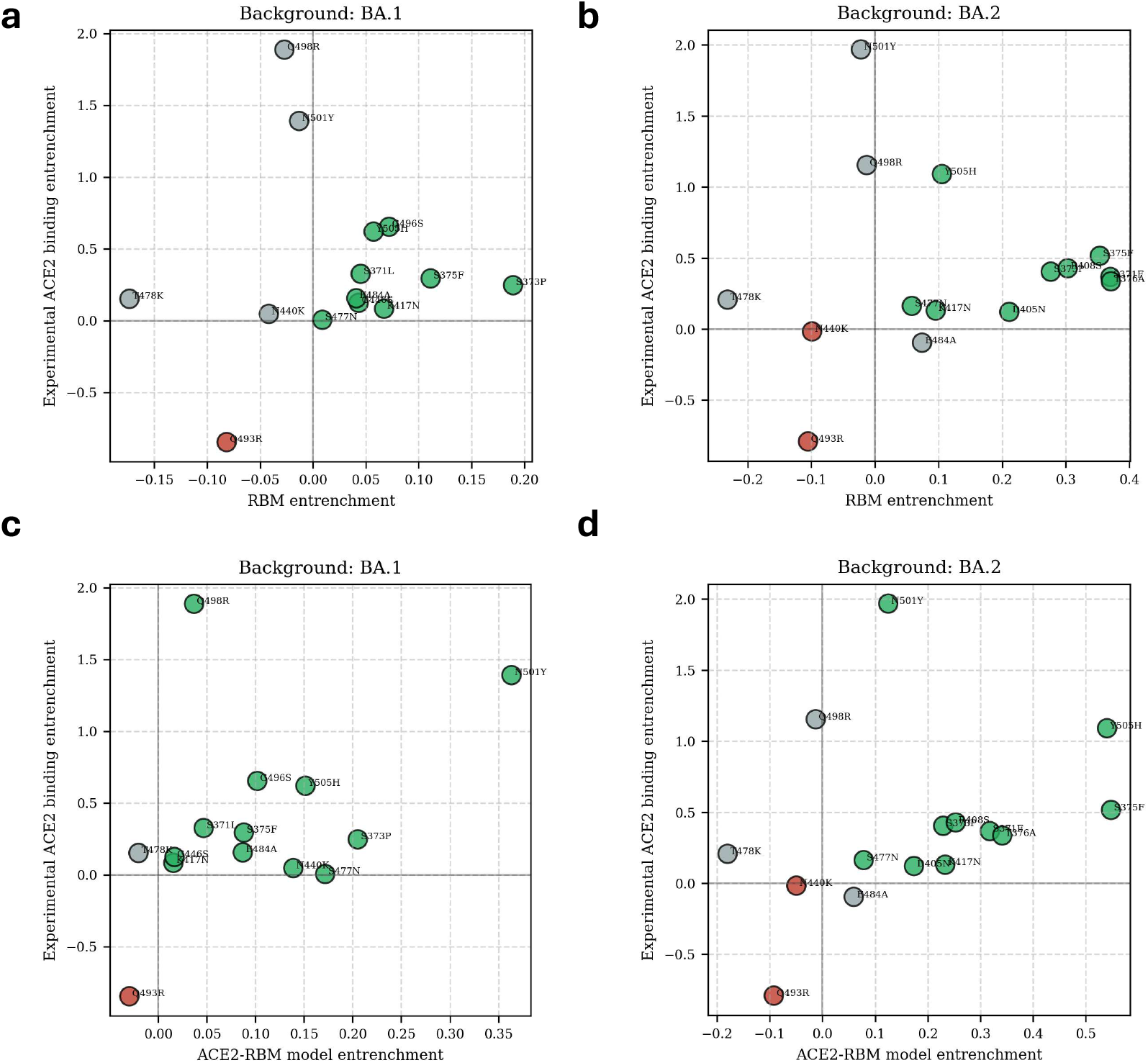
Validation of RBM+ACE2 entrenchment predictions on different backgrounds. (a-b) Mutational entrenchment in the SARS-CoV-2 Spike RBD. Comparison of mutational effects when introduced into the wild-type background versus their removal from the Omicron BA.1/BA.2 background. The shift value represents the degree of entrenchment. Results are shown for both the RBM model and experimental ACE2 binding measurements ^27^. (c-d) Same as (a–b), but using the full ACE2-RBM model (calibrated to Q2 2020 ACE2 concentrations) to compute the entrenchment scores.

**Fig. S4.**
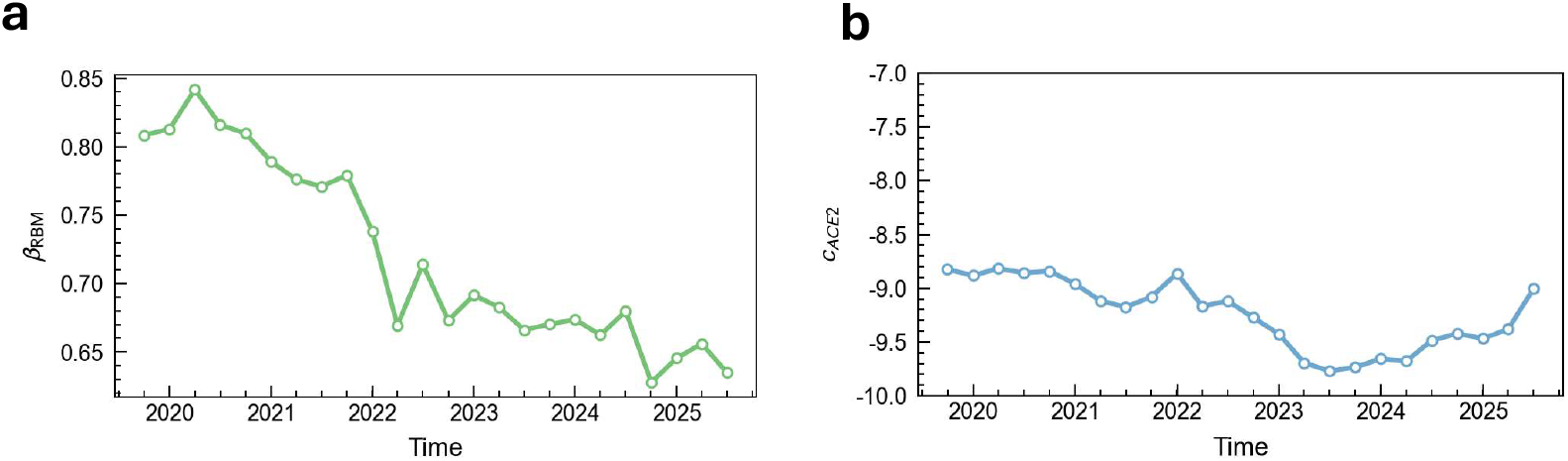
Fitted model parameters. (a) RBM inverse temperature fitted by the model as a function of time. (b) log_10_ ACE2 concentrations (mol/L) fitted by the model as a function of time.

**Fig. S5.**
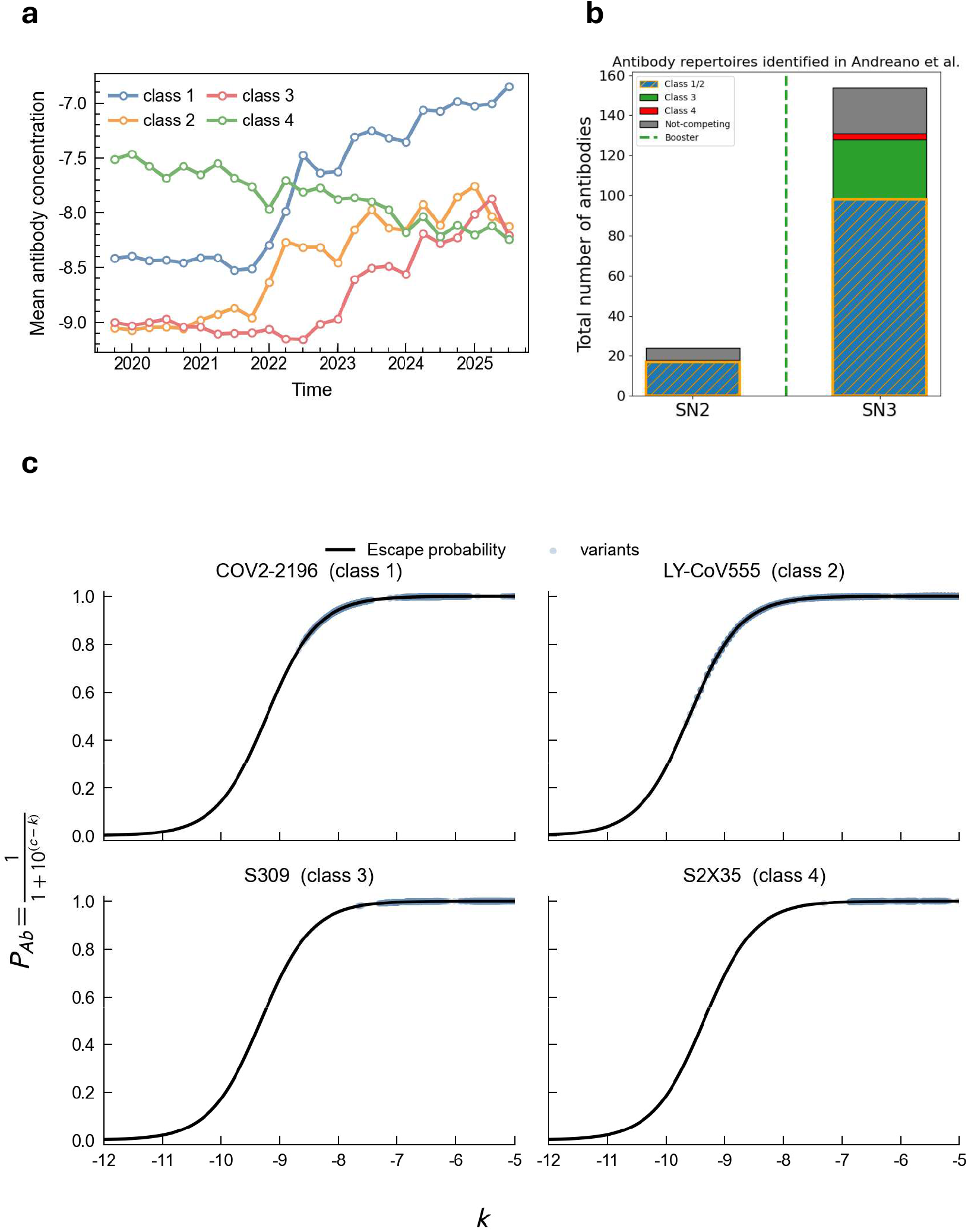
Antibody concentrations. (a) Average log_10_ Ab concentrations (g/ml) fitted by the model as a function of time by antibody class. (b) Diversity of antibodies found in serum after 2 vaccination (SN2) and 3 vaccination (SN3) doses ^28^. (c) Antibody escape probability computed for representative antibodies. Solid lines indicate the fitted model using antibody concentration *c* from the first time period, while points represent individual SARS-CoV-2 variants observed during the pandemic, positioned according to their predicted binding energy values.

**Fig. S6.**
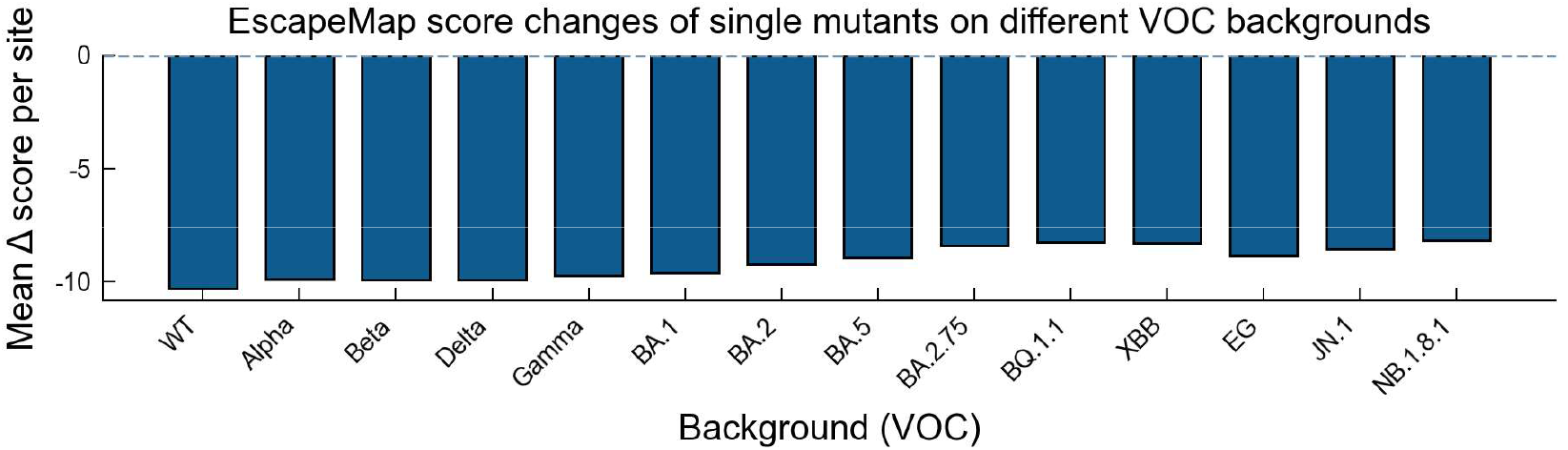
Mutational likelihood. Mean Δ score per site, defined as the difference between the EscapeMap score of the single mutant and that of the background sequence. Higher values means higher mutability. For each background, EscapeMap was fitted on period corresponding to the start of that VOC’s circulation.

**Fig. S7.**
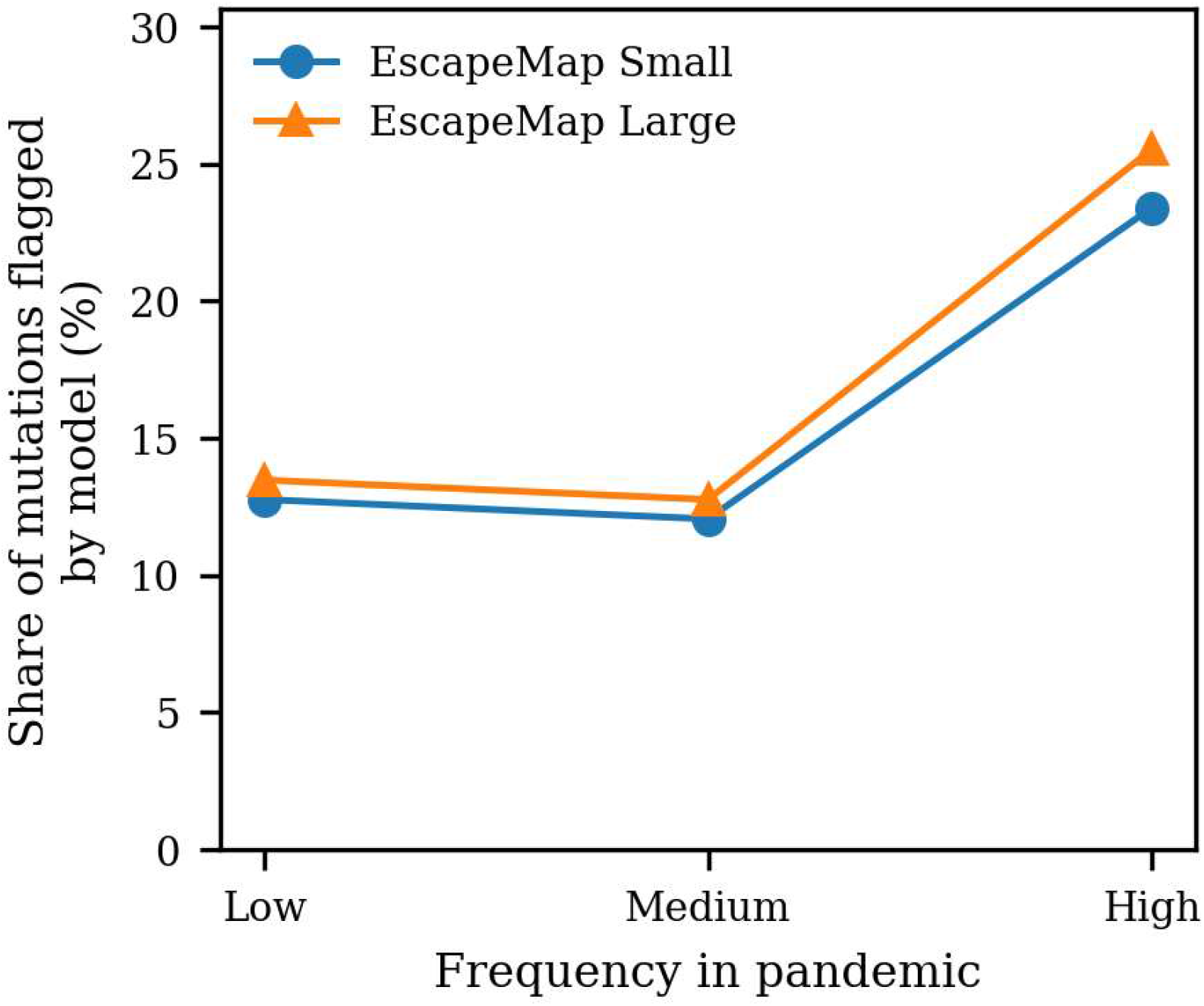
Expanding antibodies used to fit the model. Percentages of observed pandemic mutations, categorized by observed frequency during the pandemic, within model top score decile. Model was fitted on Q1 2022. Mutations are stratified into three equal-size bins based on their occurrences: Low (occurrence ≤ 284), Medium (284 *<* occurrence *<* 1443), and High (occurrence ≥ 1443). Results compare the original model, fitted with 29 antibodies identified early in the pandemic, to an expanded model incorporating 671 additional antibodies isolated from Omicron-convalescent individuals ^13,14^.

**Fig. S8.**
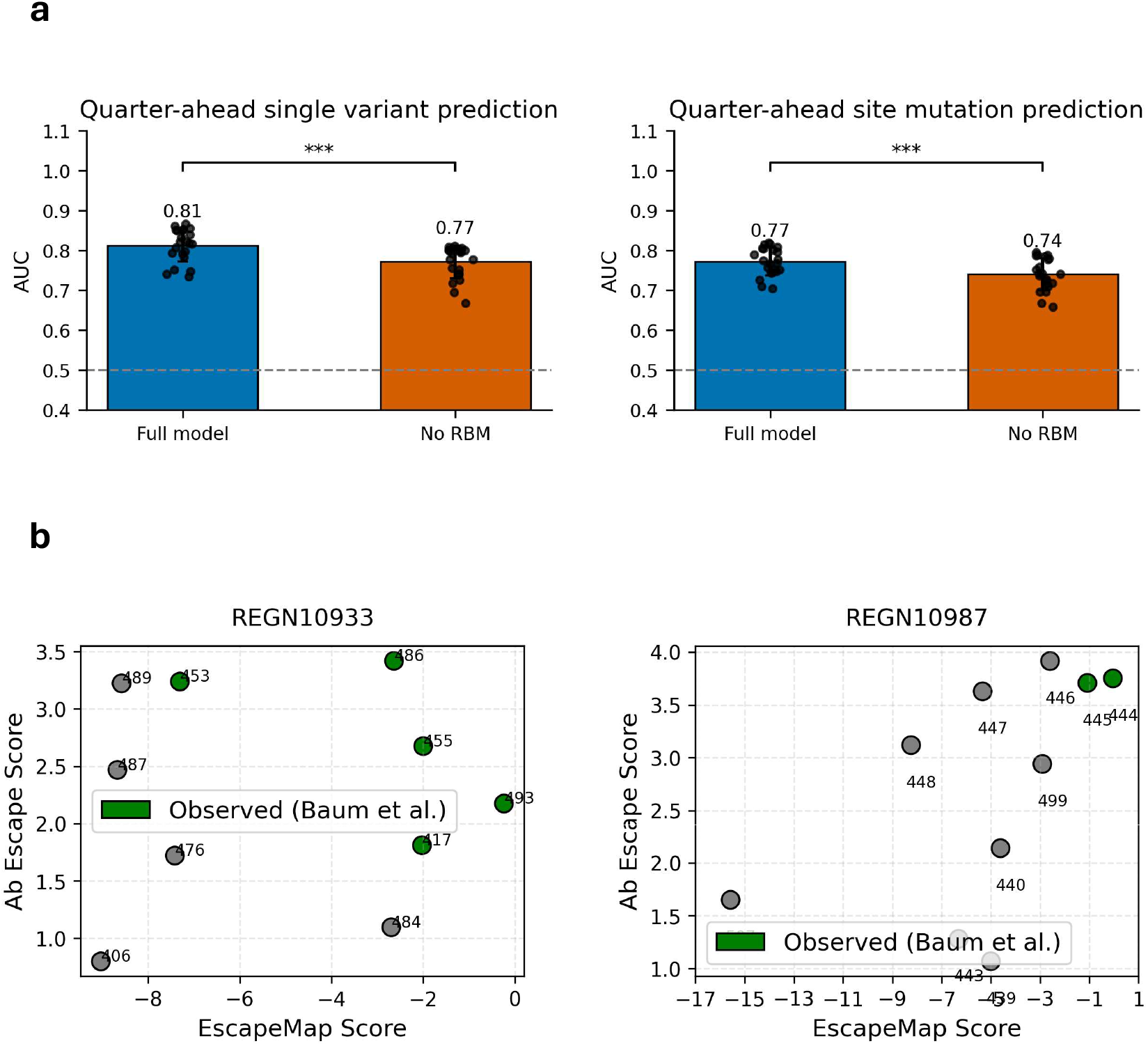
Prediction of mutated sites. (a) Predictive performance (AUC) of EscapeMap, which incorporates ACE2–Ab–RBM probabilities, compared to a baseline model that includes only ACE2–Ab terms and omits RBM-based viability, across different timepoints. AUC measures the model’s ability to correctly identify true positives against its rate of incorrectly identified false positives at all possible classification thresholds. Performance is assessed on two tasks: prediction of single-amino-acid variant effects (left) and site-level mutational tolerance (right). Asterisks indicate statistically significant differences based on paired t-tests (*p <* 0.01 for both tasks). (b) Scatter plot of site scores from EscapeMap (ACE2–Ab–RBM) (y-axis) versus antibody-only (Ab) scores (x-axis), showing the top 10 DMS-ranked escape sites. The Ab-only model reflects escape potential without viability or ACE2 constraints; a positive score indicates strong escape potential. In contrast, a positive EscapeMap score indicates that mutations at the site enable escape while maintaining viability and ACE2 binding. Observed escape sites from serial passaging ^35^ are highlighted in green. Sites 446 and 489 illustrate the model’s utility: they have good Ab-only scores but are not selected for escape, a fact captured by their low EscapeMap scores, which account for detrimental effects on protein expression and ACE2 binding, respectively.

**Fig. S9.**
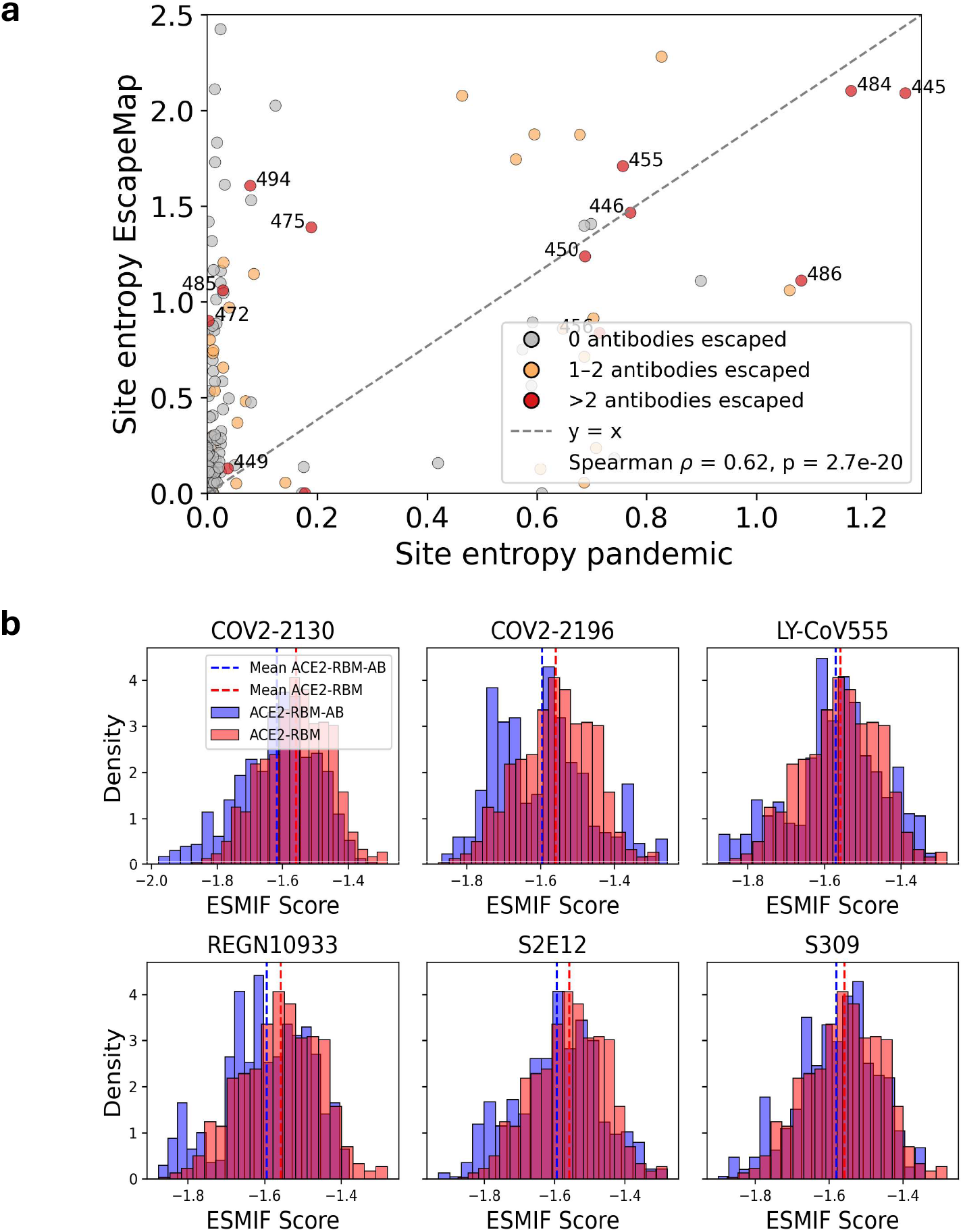
Validation of artificial sequences. (a) Comparison of site entropies of unique sequences generated using EscapeMap without antibody pressure (ACE2–RBM) and sequences found in Nextstrain. Colors indicate how many antibodies can be escaped (site max escape*>* 0.9 in DMS) when mutating that site. (b) Histograms of ESMIF scores for variants generated by EscapeMap under high antibody pressure (ACE2–Ab–RBM, log_10_ *C* = − 6 for the indicated antibody), compared to sequences generated by EscapeMap without antibody pressure (ACE2–RBM).

**Fig. S10.**
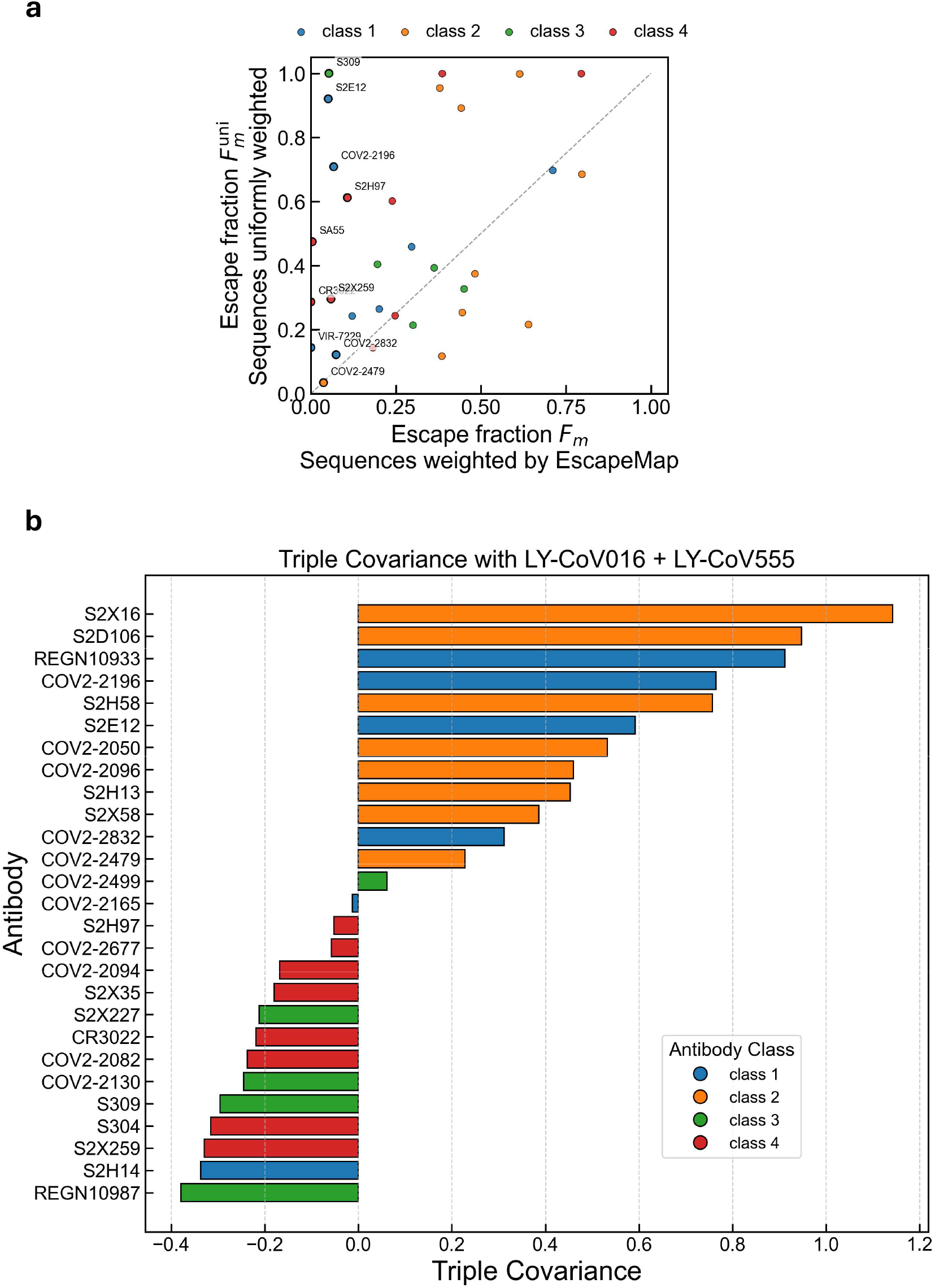
Antibody cocktails. (a) Scatter plots comparing the fraction of viable sequences predicted to escape each antibody (*F*_*m*_) with the corresponding uniform baseline 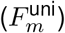. Each point represents one antibody. Lower *F*_*m*_ values indicate a reduced proportion of viable sequences capable of escape when ACE2 binding and sequence viability constraints are applied. (b) Triple covariance for 3-antibody cocktails containing LY-CoV016, LY-CoV555, and a third antibody. Classes 3 and 4 antibodies are the best candidates.

**Fig. S11.**
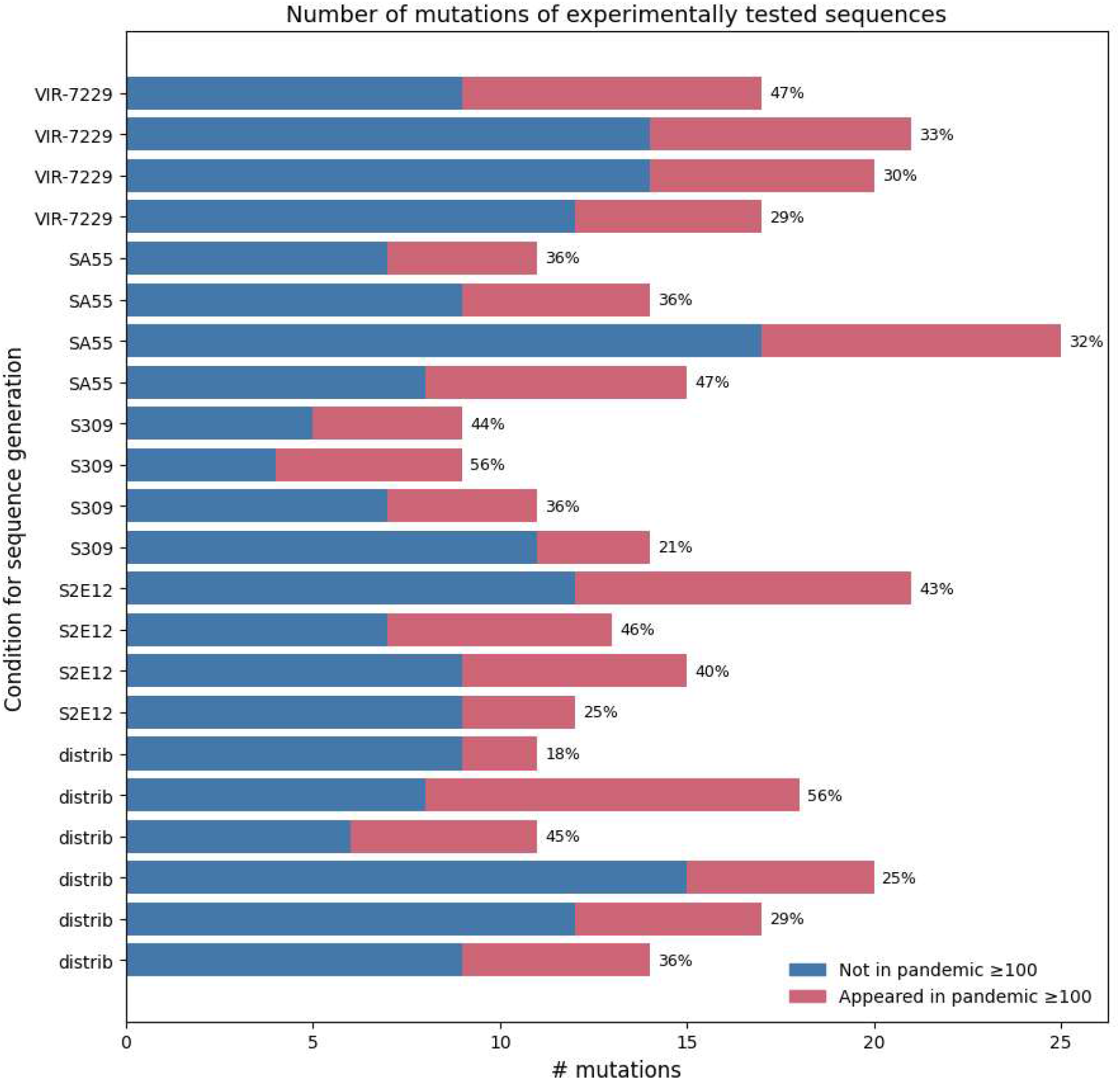
Mutational composition of experimentally generated sequences. Sequences were either generated without antibody pressure (“distrib”) or with a single antibody set to concentration log_10_ *C* = − 6. Bars show the number of mutations per sequence, split between mutations that appeared during the pandemic at least 100 times (red) and those not observed or rare (blue). Percentages indicate the fraction of mutations that were observed in the pandemic.

**Fig. S12.**
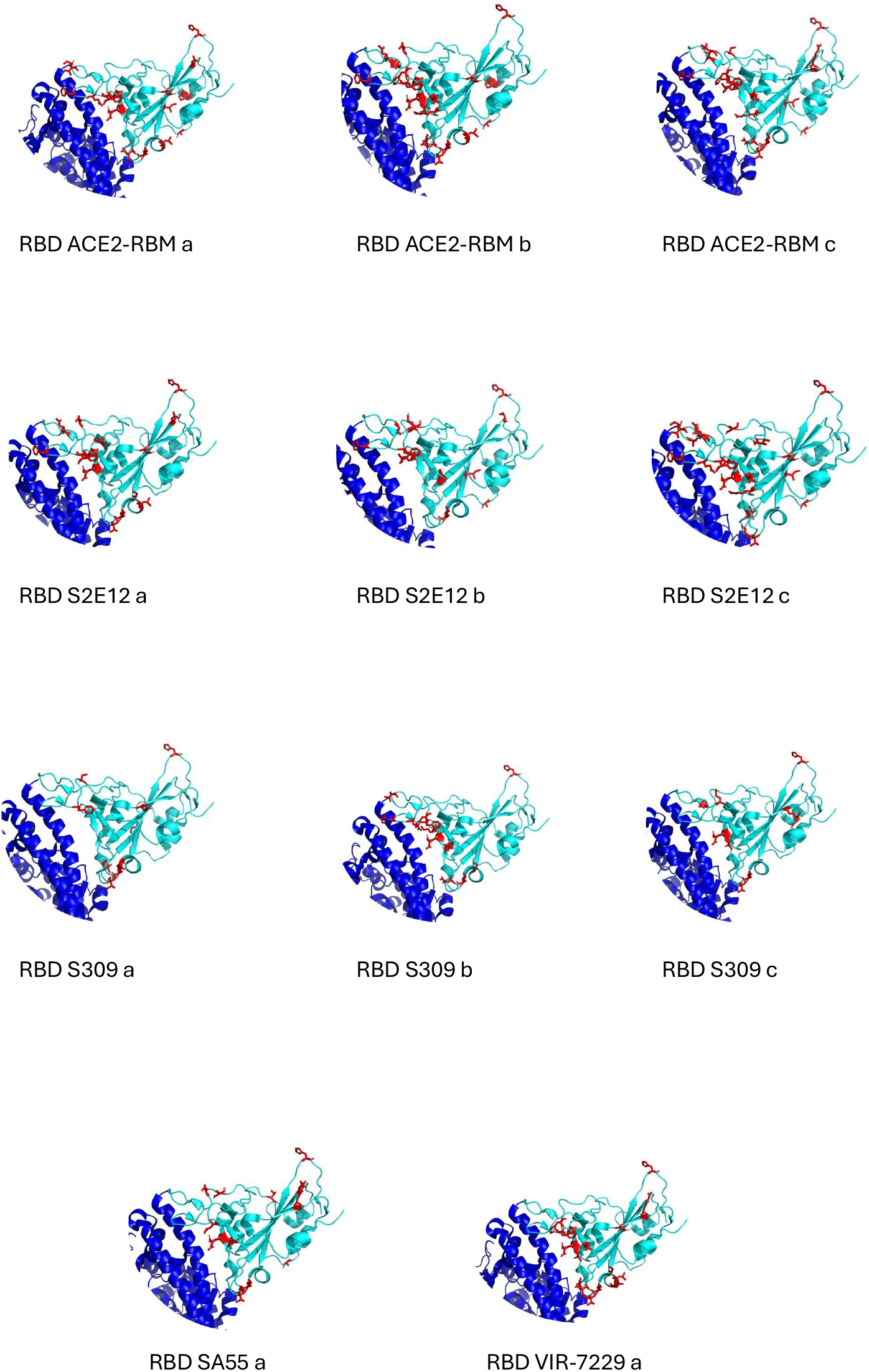
Mutated sites in expressed sequences. Structural representation of RBD, in complex with ACE2 (PDB: 6M0J), for every tested sequence that could be expressed. Red = mutations in sequence.

**Fig. S13.**
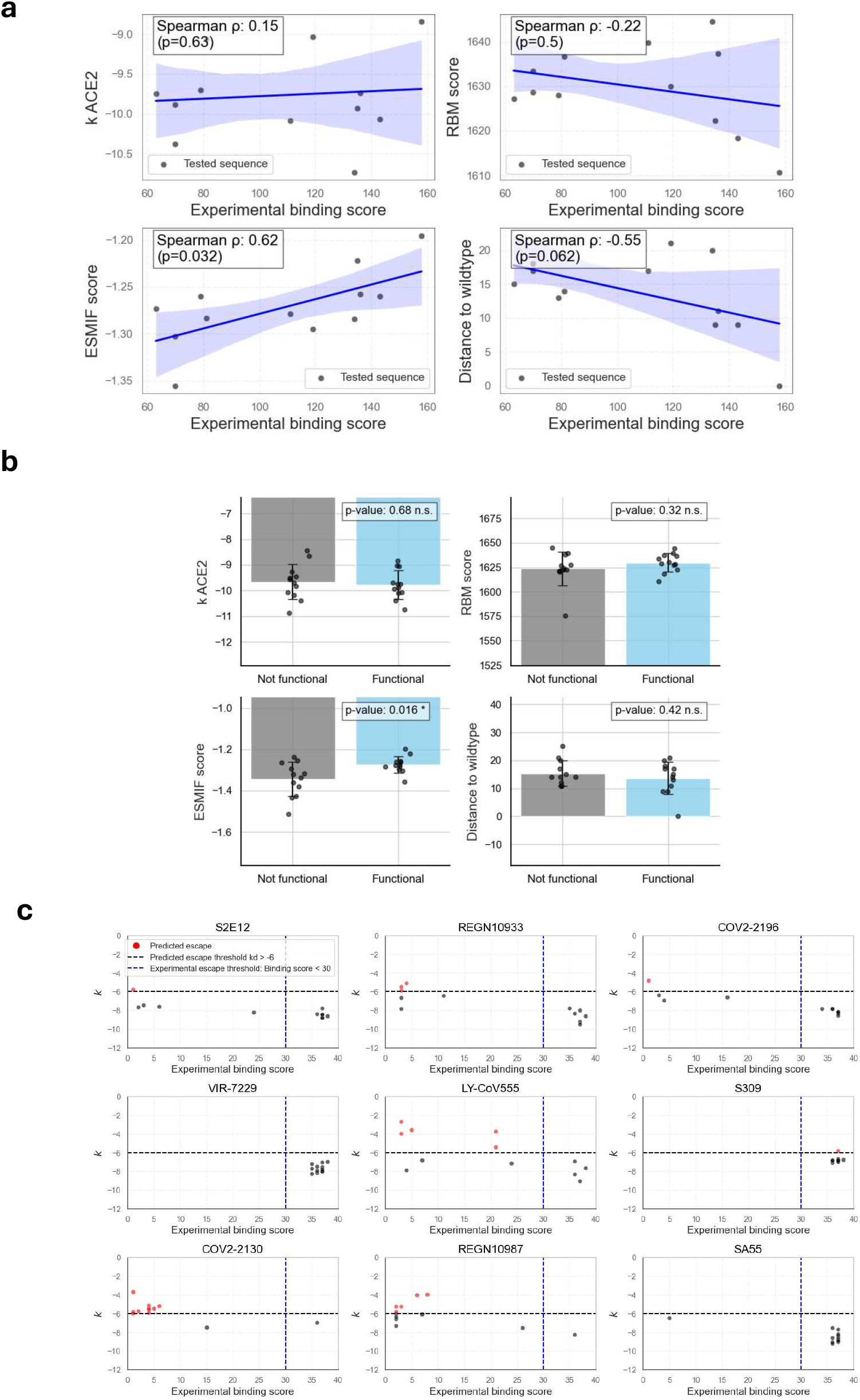
Measured binding for generated sequences. (a) Correlation between ACE2 binding measured in experiment for functional sequences and (i) distance to wildtype, (ii) RBM energy, (iii) predicted ACE2 binding energy, and (iv) ESMIF score. (b) Comparison of RBM score, predicted ACE2 binding energy, ESMIF score, and distance to wildtype between functional sequences that could be expressed and non-functional sequences. Each bar shows the mean ± standard deviation, with individual data points overlaid. Statistical significance is indicated using two-sided t-tests (*p <* 0.05: *, *p >* 0.05: ns). (c) Comparison of measured and predicted binding scores for all antibodies. Each point corresponds to a functional sequence that could be expressed.

## STAR METHODS

### METHOD DETAILS

#### Sequence data

In our study, we use multiple sequence alignments (MSAs) of 2610 sequences acquired in a previous study^3^. The sequence data in FASTA format were sourced from several databases: GISAID (release 16 May 2021), Uniref90 (December 2020 release), ViPR (downloaded in September 2020), NCBI viral genomes (downloaded in September 2020), and the MERS coronavirus database (downloaded in September 2020). As part of quality control, sequences containing nonstandard amino acids and repeated sequences were removed. Additionally, to distinguish training data from test data, any sequences with greater than 90% sequence identity to the Wuhan-Hu-1 reference were filtered out, excluding all SARS-CoV-2 sequences, including closely related sequences from nonhuman hosts.

We downloaded SARS-CoV-2 sequence data from ([Nextstrain](https://nextstrain.org/pathogens/files?filter=ncov))^49^ on August 27, 2025. All site numbering and genome structure descriptions utilize the Wuhan-Hu-1/2019 (Genbank MN908947 strain as the reference. We used the complete dataset, containing 9’369’573 sequences, to enumerate RBD amino-acid mutation counts and analyze their appearance frequencies. To avoid computational memory issues during model fitting, we utilize the representative 100,000sequence dataset provided by Nextstrain. From this subset, the model is applied to sequences spanning residues S349 to G526.

#### Background distribution of RBD sequences from pre-pandemic data

We use Restricted Boltzmann Machines^21^ to model the probability distribution over RBD sequences, which we can infer from homologous sequence data. This distribution encodes essential evolutionary, functional and structural constraints acting on RBD. A Restricted Boltzmann Machine (RBM) is a joint probabilistic model for sequences and representations. It is formally defined on a bipartite, two-layer graph. Protein sequences *v* = (*v*_1_, *v*_2_, …, *v*_*N*_ ) are displayed on the Visible layer, and representations *h* = (*h*_1_, *h*_2_, …, *h*_*M*_ ) on the Hidden layer. Each visible unit takes one out of 20 values (20 amino acids). Hidden-layer unit values *h*_*μ*_ are real. The joint probability distribution of *v, h* is:

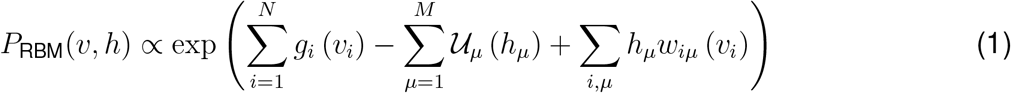

where *g*_*i*_ and *U*_*μ*_ denote, respectively, local biases and potentials acting on the visible and hidden units, and the *w*_*i,μ*_ represent the interaction weights between these layers. We use the dReLU potential 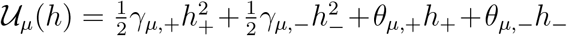, where *h*_+_ = max(*h*, 0), *h*_−_ = min(*h*, 0) . The parameters (*γ*_*μ*,+_, *γ*_*μ*,−_, *θ*_*μ*,+_, *θ*_*μ*,−_) are optimized by maximizing the product of the marginal probabilities

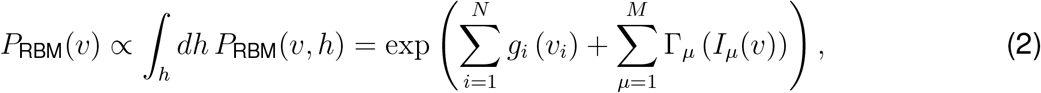

over the sequences *v* within a MSA of the protein family. The cumulant generating function 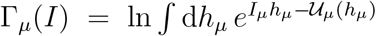, where *I*_*μ*_ = ∑_*i*_ *w*_*iμ*_ (*v*_*i*_), is computed as described in previous work^21^.

We train the RBMs using the ([PGM package](https://github.com/jertubiana/PGM))^21^ and optimize them through the Persistent Contrastive Divergence method. The number of hidden units is fixed to 100, with activation functions set to dReLU. Training is carried out employing 10 Markov chains and performing 10 Monte Carlo updates per gradient computation, for 50 iterations. To introduce regularization, we apply an L1/2 penalty on the weight parameters with a coefficient 0.12. The MSA used for training was obtained from previous work^3^ concatenated with the wildtype sequence.

### Binding to ACE2 and Antibodies

We define the probabilities that the RBD encoded by sequence *v* is bound to ACE2 and escapes antibody *i* (Ab_*i*_) through, respectively,

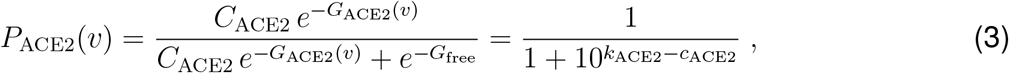

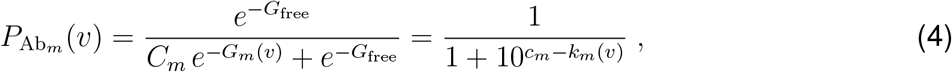

where *G*_free_, *G*_ACE2_ and *G*_*m*_ are the (dimensionless) free energies associated with the free, ACE2-bound and Ab_*m*_-bound states. We define the binding free energies *k*_*m*_(*v*) = Δ*G*_*m*_(*v*)*/* log 10 and *k*_*A*_(*v*) = Δ*G*_ACE2_(*v*)*/* log 10 for antibody *m* and ACE2, which depend on the sequence *v*. In addition, *c*_*m*_ and *c*_ACE2_ represent log10 effective concentrations of antibody *m* and ACE2 driving selection pressures over RBD. These are treated as fitted model parameters controlling immune pressure but do not represent physical concentrations within the immune system. Plots of the escape probability as a function of the binding free energy are shown, for four representative antibodies, in Figure S5.

#### Full model

Multiplying the background RBD distribution in Eq. (2) with the binding probabilities in Eqs. (3) and (4) defines the full probability distribution over the sequences *v*, with the result

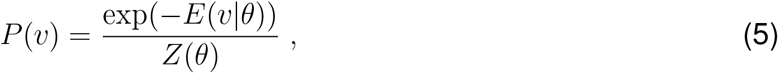

where *θ* denotes the set of all model parameters, and the effective energy *E* reads

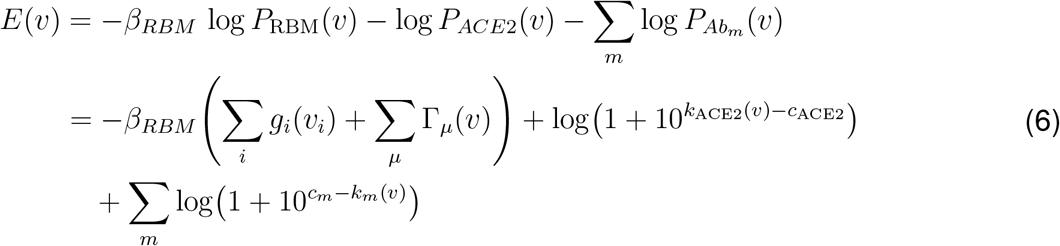

The presence of the inverse temperature parameter, *β*_*RBM*_, allows us to adjust the strength of evolutionary constraints, modulating the balance between selective pressures and sequence flexibility.

In this Boltzmann-like distribution, maximizing the total likelihood *P* (*v*) is equivalent to minimizing *E*(*v*| *θ*) + log *Z*(*θ*). Informally speaking, one wants to minimize the energy of the observed data and maximize the ones of the remaining (non observed) configurations. Accordingly, the energy term of ACE2 binding and antibody escape ensure that sequences exhibiting high receptor affinity (low *k*_ACE2_) and high escape potential (high *k*_*m*_) are assigned lower energy values, thereby increasing their statistical weight in the generative process.

Finally, we can increase the strength of selection by raising the overall model inverse temperature *β*, which amplifies the contribution of the effective energy *E*(*v*) to the sampling distribution. Accordingly, we sample sequences from the Boltzmann-like distribution

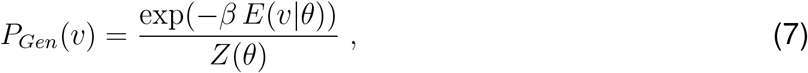

where the partition function *Z*(*θ*) ensures normalization and is defined as

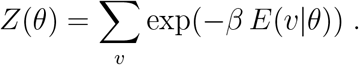

To sample from *P*_*Gen*_(*v*), we utilize a Markov Chain Monte Carlo (MCMC) approach based on the Metropolis-Hastings algorithm. This framework is particularly suitable because the partition function *Z*(*θ*), which is computationally intractable due to the vastness of the sequence space, cancels out during the calculation of the acceptance ratio. Starting from an initial sequence *v*, a mutation is proposed to generate a candidate sequence *v*^*′*^. The move is accepted with probability *α* = min(1, exp( −*β*[*E*(*v*| ^*′*^ *θ*) −*E*(*v* | *θ*)])); otherwise, the chain remains at *v*. Through repeated iterations, the system converges to a stationary distribution, allowing for the generation of sequences that accurately represent the high-fitness landscape.

#### Prediction of binding free energies for multiple variants

To model the effect of single mutations on binding free energies, we use deep mutational scans for ACE2^22^, SA55 and VIR-7229^33^, as well as data provided in [bloomlab repository](https://jbloomlab.github.io/SARS2_RBD_Ab_escape_maps/), that integrates data from several previous studies^6–12^. DMS data provided escape ratios *ϵ*_*i*_ for nearly all single mutants *i* on a Wuhan background, to all 29 antibodies. We set wildtype escape to a minimum value of 0.01, close to minimum measurable. We then define the site specific contribution as 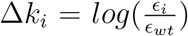. To extrapolate to variants with multiple mutations, we use an additive model in terms of site-specific contributions Δ*k*_*i*_ to compute the binding free energy *k* of sequence *v* = (*v*_1_, …, *v*_*N*_ ) with amino acids *v*_*i*_:

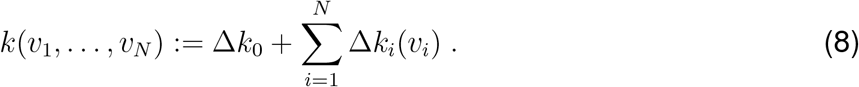

In this equation, we choose 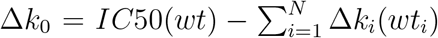 so that predicted binding free energy of wildtype matches its IC50. In practice, we compute the binding free energy using a ridge regression (L2 penalty *λ* = 0.1) on the one hot encoding of sequence amino acids. A unique property of ridge regression applied to one-hot encoding is that the predicted mutational effect for unknown amino acids is the average known mutational effect of mutations on the same site^26^. This mechanism allows the model to effectively generalize to unknown amino acids at specific positions from knowledge of the average impact of mutations on the dissociation constant. This is necessary as deep mutational scans may not cover the entirety of possible single variants.

### Experimental IC50

Fitting the linear model requires the effects of single mutations and the wildtype IC50, which defines the model’s fixed y-intercept. IC_50_ values for the 29 antibodies against the wild-type RBD were taken from^12^ for S304, S2H14, S2H97, S2H13, S2X35, S309, S2X58, S2X227, S2X16, S2H58, S2D106 and S2E12;^50^ for COV2-2677, COV2-2165, COV2-2082, COV2-2094, COV2-2130, COV2-2050, COV2-2832, COV2-2096, COV2-2479, COV2-2499 and COV2-2196;^11^ for REGN10987, REGN10933 and LY-CoV016;^51^ for LY-CoV555;^9^ for S2X259; and^6^ for CR3022.

We also source IC50 data for two larger antibody cohorts^13,14^. The first set, used in the expanded version of the model, comprises 671 antibodies from BA.1 convalescent individuals. The second set is used to test the model’s predictive capacity for BA.1; for this, we extract wildtype and BA.1 IC50 values for 438 antibodies identified in WT convalescents.

#### Fitting ACE2-AB-RBM model on pandemic data

We want to find the parameter set *θ* that maximizes the model average log-likelihood ⟨log *P* (*s*) _datase_⟩of the sequences in the training set, which includes pandemic data. For simplicity, we collectively denote as *θ* the set of parameters encompassing antibody and ACE2 log-concentrations {*c*_*i*_} and the inverse temperature *β*_*RBM*_ of the RBM component.

The gradient of the log-likelihood is given by

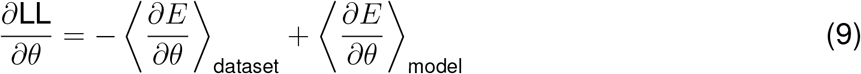

In practice, we choose a minibatch with a single sequence, randomly drawn from the dataset at each step of the learning algorithm. We then sample another sequence from the model distribution *P*, and compute the energy difference between these two sequences. Using PyTorch’s autodifferentiation tools we evaluate the gradient. Last of all, we perform a gradient update step of *θ* based on this computation.

To sample from the model distribution, the conventional method involves performing a complete Markov Chain Monte Carlo (MCMC) simulation for each gradient step, continuing until equilibrium is reached. A more computationally efficient alternative is Persistent Contrastive Divergence (PCD), a technique introduced by Tieleman for training Restricted Boltzmann Machines^52^. Instead of initializing each MCMC run from scratch, PCD reuses samples from the previous iteration, updating them with a limited number of MCMC steps before the next gradient evaluation. When the model parameters change gradually, this approach effectively maintains equilibrium, providing accuracy comparable to standard MCMC sampling while reducing computational cost.

#### RBM site mutational score

For each site *i*, we define the RBM log-likelihood score of mutating site *i* in the wild-type background:

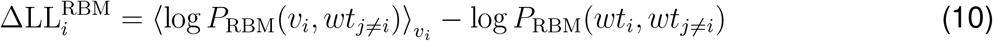

where 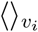 denotes the average over the set of all single mutants with amino-acid *v*_*i*_ at site *i* .

#### Fraction of escaping variants

We compute the escape fraction *F*_*m*_ for a specific antibody, *Ab*_*m*_, by integrating its density of variants having binding free energies *f*_*m*_(*k*_*m*_) above an escape threshold *k*_*esc*_:

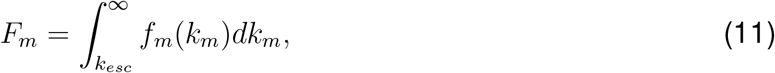

where *f*_*m*_(*k*_*m*_) *dk*_*m*_ is the fraction of viable variants whose binding free energy for antibody *m* is contained in the range [*k*_*m*_, *k*_*m*_ + *dk*_*m*_]:

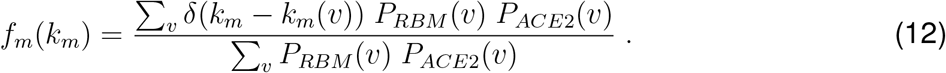

The *F*_*m*_ score represents the probability that a randomly selected functional sequence escapes the given antibody, using an escape threshold of *k*_*esc*_ = −6.

This approach can be extended to evaluate antibody cocktails by analyzing the joint probability distribution of binding free energies across all antibodies in the set. The total escape fraction for the cocktail represents the probability that a randomly selected functional variant can simultaneously escape every antibody present. This value is determined by accumulating the probability of all variants whose binding free energy exceeds the defined escape threshold for each and every antibody in the cocktail.

#### Antibody synergy

To quantify antibody synergy, we compute the covariance between the binding free-energies of viable RBD sequences to two antibodies, say, *m* and *m*^*′*^:

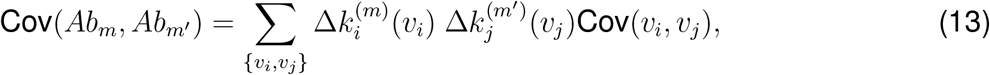

where Cov(*v*_*i*_, *v*_*j*_) represents the covariance of amino-acid occurrences when sampling sequences from the distribution *P* (*v*), and 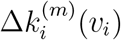 denotes the effect on the binding free energy for antibody *m* of the amino acid *v* on site *i*.

Synergetic pairs of antibodies cannot be escaped by the same mutation *v*_*i*_, requiring 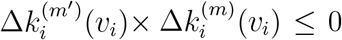. This statement extends to positively correlated mutations (*v*_*i*_, *v*_*j*_), *i*.*e*. such that Cov(*v*_*i*_, *v*_*j*_) *>* 0: such mutations should have opposite effects on the two binding free energies, 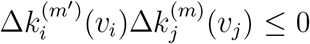. Finally, pairs of mutations (*v*_*i*_, *v*_*j*_) that would increase both binding free energies and help antibody escape, *i*.*e*. such that 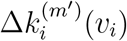 and 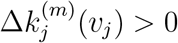, should be disfavored and exhibit zero or negative covariance, Cov(*v*_*i*_, *v*_*j*_) ≤ 0.

This formalism extends to cocktails of k antibodies by generalizing the covariance of aminoacid occurrences with the k-order centered moment. For cocktails of 3 antibodies, we obtain:

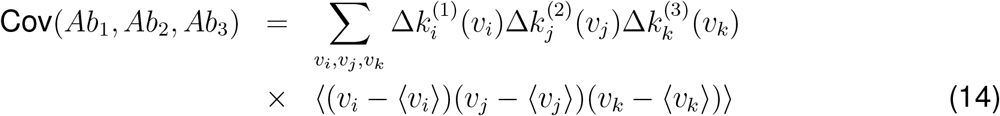

where the sum runs over all triplets of mutations (*v*_*i*_, *v*_*j*_, *v*_*k*_).

#### Sequence Generation under Different Selective Pressures

For each experimental condition, we generate a batch of 1,000 RBD sequences by sampling from the gap-less distribution *P* (*v*) (Eq. 5). Samples are obtained using Markov Chain Monte Carlo (MCMC, see Methods).

To simulate immune pressure, sequences are drawn from the full model (Eq. 5), which includes the ACE2-RBM energy landscape and the antibody term favoring escape from a given antibody with antibody concentration parameters set to log_10_ *c* = −6.

As controls, we also sample from perturbed distributions. To test the effect of reducing structural constraints, we increase the RBM temperature (*β*_RBM_ ∈ {0.1, 0.2} ) to relax evolutionary penalties. Conversely, to test reduced ACE2 affinity, we use a higher ACE2 concentration (log_10_ *C*_ACE2_ ∈ {−6, −4}), diminishing the selection for tight receptor binding.

#### Site entropy

To quantify the mutational variability at each site *i* of the RBD, we compute the Shannon entropy:

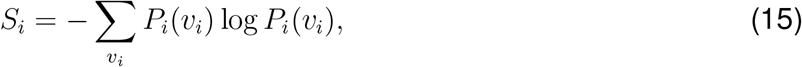

where *P*_*i*_(*v*_*i*_) is the marginal probability of observing amino acid *v*_*i*_ at position *i*. High entropy indicates that the site tolerates diverse mutations, while low entropy reflects evolutionary constraints or functional importance.

#### Sequence scoring with ESM Inverse Folding

To evaluate sequence likelihoods by a state of the art machine-learning method learned on structures, we use *ESM Inverse Folding* available in [ESMIF repository](https://github.com/facebookresearch/esm/tree/main/examples/inverse_folding), which assigns a log-likelihood score to each sequence given a protein structure^26^. In this approach, the structural context is provided by the PDB file, while the FASTA file contains the sequences to be assessed. The loglikelihood scores quantify how well each sequence aligns with the given structure. For ACE2, we conditioned binding on pdb 6M0J, for LY-CoV555^53^ we used pdb 7KMG, for LY-COV016 we used pdb 7C01, for REGN10987 we used pdb 9LYP.

#### Recombinant RBD proteins and monoclonal antibodies

Codon-optimized nucleotide fragments encoding the RBD proteins followed by C-terminal tags (Hisx8-tag and AviTag) were synthesized and cloned into pcDNA3.1/Zeo(+) expression vector (Thermo Fisher Scientific). A single point mutation, G504E, was introduced into the RBD-encoding constructs using the QuickChange Site-Directed Mutagenesis kit (Agilent Technologies) following the manufacturer’s instructions. Human angiotensin-converting enzyme 2 (ACE2) ectodomain and RBD proteins were produced by transient transfection of exponentially growing Freestyle 293-F suspension cells (Thermo Fisher Scientific) using polyethylenimine (PEI) precipitation method, purified from culture supernatants by high-performance chromatography using the Ni Sepharose® Excel Resin according to manufacturer’s instructions (Citivia), dialyzed against PBS using Slide-A-Lyzer® dialysis cassettes (Thermo Fisher Scientific), quantified using NanoDrop 2000 instrument (Thermo Fisher Scientific), and controlled for purity by SDS-PAGE using NuPAGE 3-8% Tris-acetate gels (Life Technologies) as previously described^54^. Purified AviTagged-ACE2 protein was biotinylated using the Enzymatic Protein Biotinylation Kit following the manufacturer’s procedures (Sigma-Aldrich). Human anti-SARS-CoV2 RBD and negative control (mGO53) IgG1 antibodies^55^ were produced by transient co-transfection of Freestyle™ 293-F suspension cells (Thermo Fisher Scientific), purified by affinity chromatography using Protein G Sepharose® 4 Fast Flow (GE Healthcare) according to manufacturer’s instructions (Citivia), and quantified using NanoDrop 2000 instrument (Thermo Fisher Scientific).

### ELISA

ELISAs were performed as previously described^54^. Briefly, high-binding 96-well ELISA plates (Costar, Corning) were coated overnight with 250 ng/well of purified recombinant RBD proteins. After washings with 0.05% Tween 20-PBS (washing buffer), plates were blocked 2 h with 2% BSA, 1 mM EDTA, 0.05% Tween 20-PBS (Blocking buffer), washed, and incubated with serially diluted purified IgG mAbs in PBS. Recombinant IgG1 antibodies were tested at 10 μg/ml, and 7 consecutive 1:4 dilutions in PBS. After washings, plates were revealed by incubation for 1 h with goat HRP-conjugated anti-human IgG (Jackson ImmunoReseach, 0.8 μg/ml final) and by adding 100 μl of HRP chromogenic substrate (ABTS solution, Euromedex) after washing steps. Optical densities were measured at 405 nm (OD405nm) after 1 h incubation at room temperature, and background values given by incubation of PBS alone in coated wells were subtracted. For ACE2 binding to immobilized RBD proteins, plates were incubated with recombinant biotinylated ACE2 ectodomain at 50 μg/ml and 7 consecutive 1:2 dilutions. After washings, plates were revealed by incubation for 30 min with streptavidin HRP-conjugated (BD Biosciences) as described above. Experiments were performed using HydroSpeed™ microplate washer and Sunrise™ microplate absorbance reader (Tecan Männedorf, Switzerland). Area under the curve (AUC) values were calculated from ELISA titration curves obtained in duplicate.

**Table 2:**
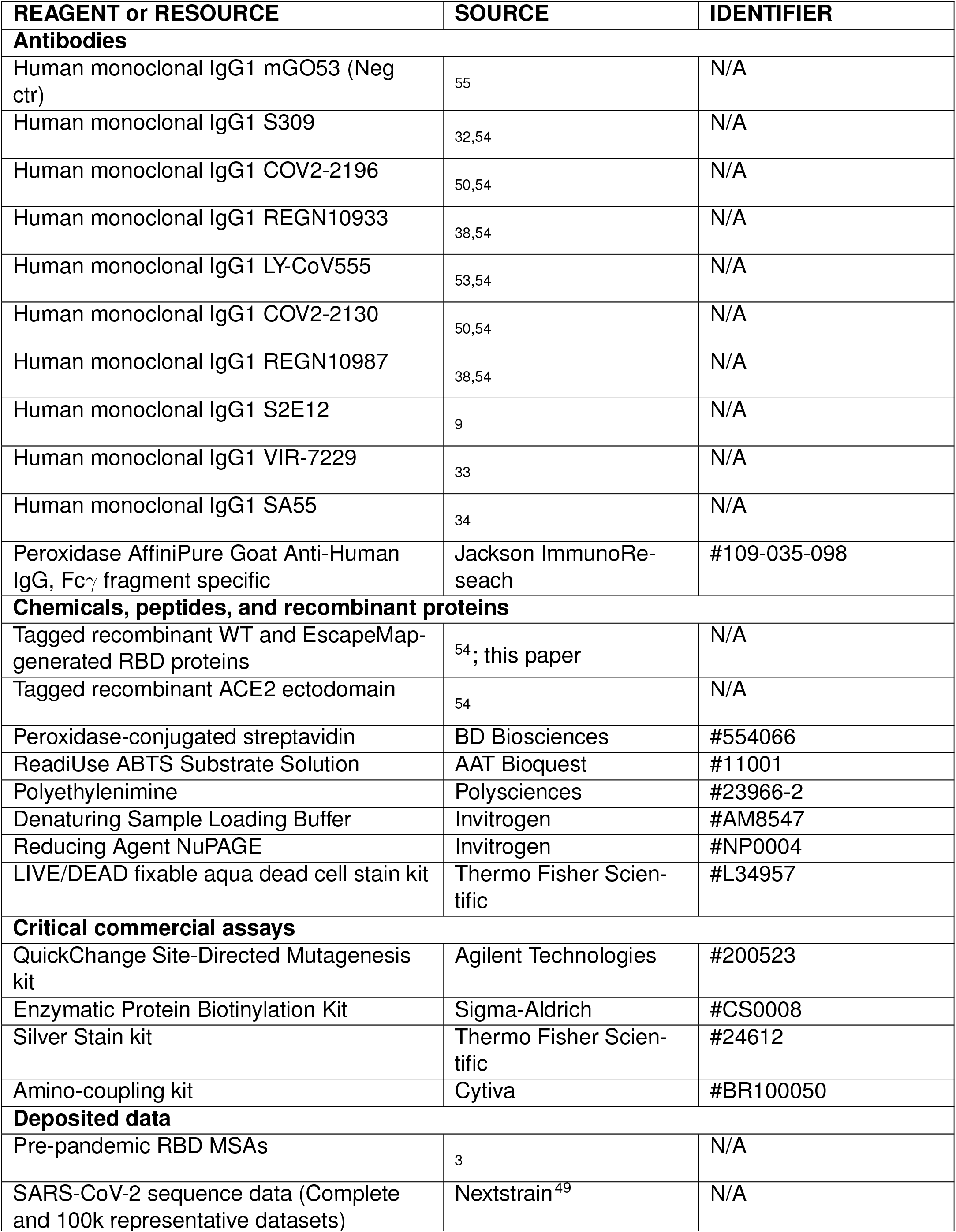
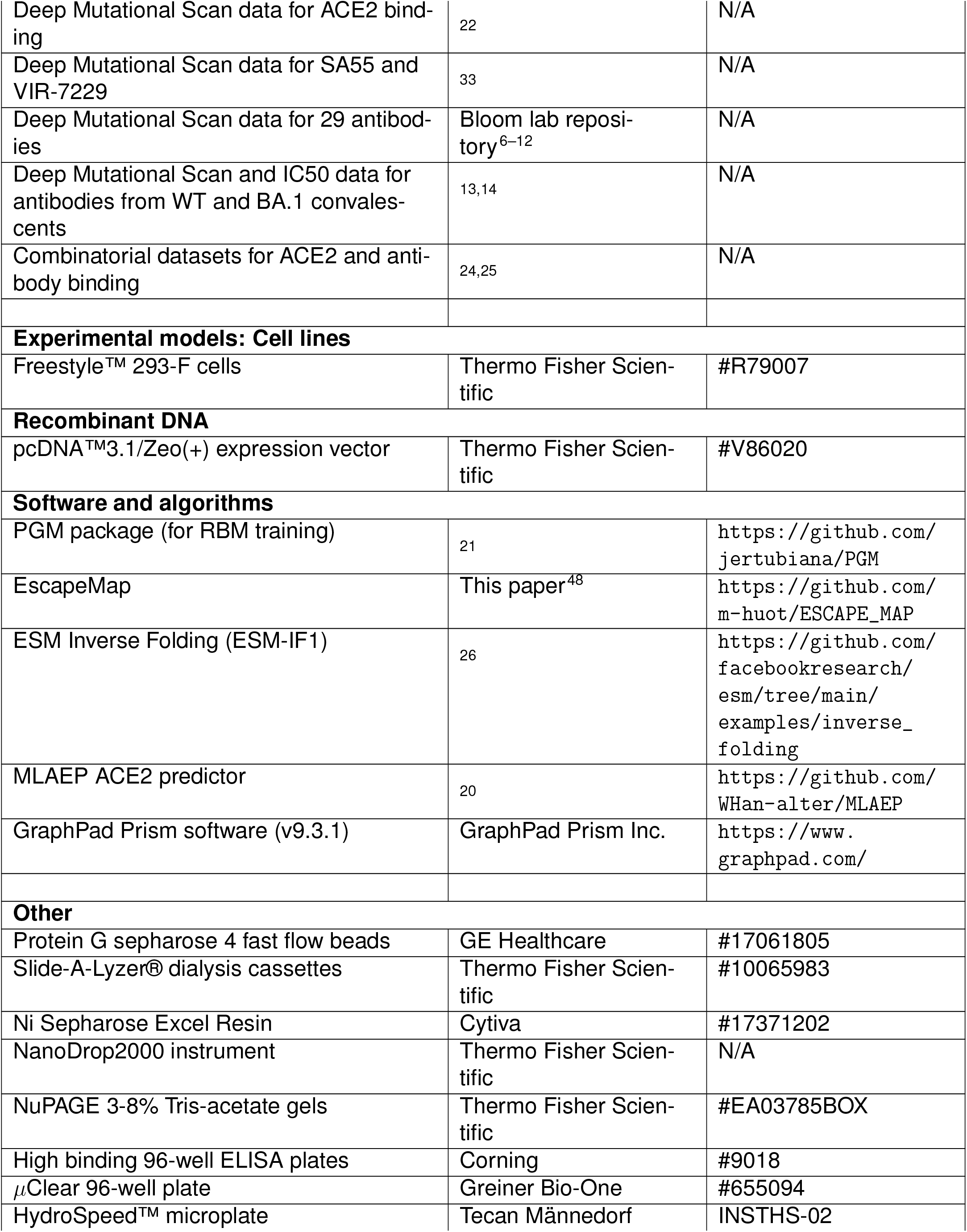
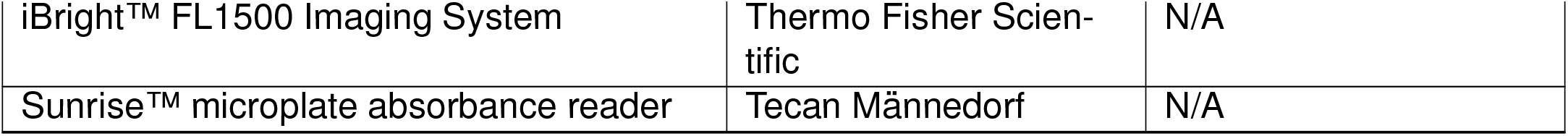
Key Resources Table

| REAGENT or RESOURCE | SOURCE | IDENTIFIER |
| --- | --- | --- |
| **Antibodies** |  |  |
| Human monoclonal IgG1 mGO53 (Neg ctr) | 55 | N/A |
| Human monoclonal IgG1 S309 | 32,54 | N/A |
| Human monoclonal IgG1 COV2-2196 | 50,54 | N/A |
| Human monoclonal IgG1 REGN10933 | 38,54 | N/A |
| Human monoclonal IgG1 LY-CoV555 | 53,54 | N/A |
| Human monoclonal IgG1 COV2-2130 | 50,54 | N/A |
| Human monoclonal IgG1 REGN10987 | 38,54 | N/A |
| Human monoclonal IgG1 S2E12 | 9 | N/A |
| Human monoclonal IgG1 VIR-7229 | 33 | N/A |
| Human monoclonal IgG1 SA55 | 34 | N/A |
| Peroxidase AffiniPure Goat Anti-Human IgG, Fcγ fragment specific | Jackson ImmunoResearch | #109-035-098 |
| **Chemicals, peptides, and recombinant proteins** |  |  |
| Tagged recombinant WT and EscapeMap-generated RBD proteins | <sup>54</sup>; this paper | N/A |
| Tagged recombinant ACE2 ectodomain | 54 | N/A |
| Peroxidase-conjugated streptavidin | BD Biosciences | #554066 |
| ReadiUse ABTS Substrate Solution | AAT Bioquest | #11001 |
| Polyethylenimine | Polysciences | #23966-2 |
| Denaturing Sample Loading Buffer | Invitrogen | #AM8547 |
| Reducing Agent NuPAGE | Invitrogen | #NP0004 |
| LIVE/DEAD fixable aqua dead cell stain kit | Thermo Fisher Scientific | #L34957 |
| **Critical commercial assays** |  |  |
| QuickChange Site-Directed Mutagenesis kit | Agilent Technologies | #200523 |
| Enzymatic Protein Biotinylation Kit | Sigma-Aldrich | #CS0008 |
| Silver Stain kit | Thermo Fisher Scientific | #24612 |
| Amino-coupling kit | Cytiva | #BR100050 |
| **Deposited data** |  |  |
| Pre-pandemic RBD MSAs | 3 | N/A |
| SARS-CoV-2 sequence data (Complete and 100k representative datasets) | Nextstrain<sup>49</sup> | N/A |
| *Continued on next page...* |  |  |
| Deep Mutational Scan data for ACE2 binding | 22 | N/A |
| Deep Mutational Scan data for SA55 and VIR-7229 | 33 | N/A |
| Deep Mutational Scan data for 29 antibodies | Bloom lab repository<sup>6–12</sup> | N/A |
| Deep Mutational Scan and IC50 data for antibodies from WT and BA.1 convalescents | 13,14 | N/A |
| Combinatorial datasets for ACE2 and antibody binding | 24,25 | N/A |
| **Experimental models: Cell lines** |  |  |
| Freestyle™ 293-F cells | Thermo Fisher Scientific | #R79007 |
| **Recombinant DNA** |  |  |
| pcDNA™3.1/Zeo(+) expression vector | Thermo Fisher Scientific | #V86020 |
| **Software and algorithms** |  |  |
| PGM package (for RBM training) | 21 | https://github.com/jertubiana/PGM |
| EscapeMap | This paper<sup>48</sup> | https://github.com/m-huot/ESCAPE_MAP |
| ESM Inverse Folding (ESM-IF1) | 26 | https://github.com/facebookresearch/esm/tree/main/examples/inverse_folding |
| MLAEP ACE2 predictor | 20 | https://github.com/WHan-alter/MLAEP |
| GraphPad Prism software (v9.3.1) | GraphPad Prism Inc. | https://www.graphpad.com/ |
| **Other** |  |  |
| Protein G sepharose 4 fast flow beads | GE Healthcare | #17061805 |
| Slide-A-Lyzer® dialysis cassettes | Thermo Fisher Scientific | #10065983 |
| Ni Sepharose Excel Resin | Cytiva | #17371202 |
| NanoDrop2000 instrument | Thermo Fisher Scientific | N/A |
| NuPAGE 3-8% Tris-acetate gels | Thermo Fisher Scientific | #EA03785BOX |
| High binding 96-well ELISA plates | Corning | #9018 |
| μClear 96-well plate | Greiner Bio-One | #655094 |
| HydroSpeed™ microplate | Tecan Männedorf | INSTHS-02 |
| Continued on next page... |  |  |

*...continued from previous page*

| **REAGENT or RESOURCE** | **SOURCE** | **IDENTIFIER** |
| --- | --- | --- |
| iBright™ FL1500 Imaging System | Thermo Fisher Scientific | N/A |
| Sunrise™ microplate absorbance reader | Tecan Männedorf | N/A |

